# Distinct molecular pathways govern presynaptic homeostatic plasticity

**DOI:** 10.1101/2020.12.21.423841

**Authors:** Anu G. Nair, Paola Muttathukunnel, Martin Müller

**Affiliations:** Department of Molecular Life Sciences, University of Zurich, Winterthurerstrasse 190, 8057 Zurich, Switzerland; Department of Neuroscience, Karolinska Institutet, 17177 Stockholm, Sweden; Neuroscience Center Zurich, University of Zurich/ETH Zurich, 8057 Zurich, Switzerland

## Abstract

Presynaptic homeostatic plasticity (PHP) stabilizes synaptic transmission by counteracting impaired neurotransmitter receptor function through increased presynaptic release. PHP is thought to be triggered by impaired receptor function, and to involve a stereotypic signaling pathway. However, here we demonstrate that different receptor perturbations that similarly reduce synaptic activity result in vastly different responses at the *Drosophila* neuromuscular junction. While receptor inhibition by the glutamate receptor (GluR) antagonist γ-DGG is not compensated by PHP, the GluR inhibitors PhTx-433 and Gyki induce compensatory PHP. Intriguingly, PHP triggered by PhTx and Gyki involve separable signaling pathways, including inhibition of distinct GluR subtypes, differential modulation of the active-zone scaffold Bruchpilot, and short-term plasticity. Moreover, while PHP upon Gyki treatment does not require genes promoting PhTx-induced PHP, it involves presynaptic Protein Kinase D. Thus, synapses not only respond differentially to similar activity impairments, but achieve homeostatic compensation via distinct mechanisms, highlighting the diversity of homeostatic signaling.

## Introduction

A variety of homeostatic signaling systems stabilize neural function by counteracting diverse perturbations (Delvendahl et al., 2019; Frank et al., 2006; Ibata et al., 2008; Keck et al., 2013; Petersen et al., 1997; Teichert et al., 2017; Turrigiano et al., 1998). Chemical synapses compensate for neural activity perturbations through homeostatic regulation of neurotransmitter release (Frank et al., 2006; Petersen et al., 1997) or neurotransmitter receptors (Turrigiano et al., 1998). There are also emerging links between homeostatic synaptic plasticity and neural disease, such as autism spectrum disorders, schizophrenia, or amyotrophic lateral sclerosis (Genç et al., 2020; Perry et al., 2017; Tatavarty et al., 2020).

Presynaptic homeostatic plasticity (PHP) is a major form of homeostatic plasticity that is characterized by homeostatic upregulation of presynaptic release in response to reduced postsynaptic neurotransmitter receptor activity (Delvendahl and Müller, 2019). PHP has been observed at diverse synapses in different species (Delvendahl et al., 2019; Frank et al., 2006; Wang et al., 2018), implying evolutionary conservation. Experimentally, PHP is induced by pharmacological or genetic neurotransmitter receptor impairment (Frank et al., 2006; Petersen et al., 1997). Correspondingly, reduced postsynaptic excitation caused by receptor impairment, as quantified by a reduction in quantal size, is thought to trigger homeostatic signaling. In this regard, Ca^2+^ flux through postsynaptic receptors has been hypothesized to be a major synaptic variable sensed by the signaling system promoting PHP (Frank et al., 2006; Ouanounou et al., 2016; Wang et al., 2016). In agreement, postsynaptic signaling by CaMKII and the Ca^2+^-binding protein Peflin has been implicated in PHP at the *Drosophila* neuromuscular junction (NMJ) (Haghighi et al., 2003; Kikuma et al., 2019; Li et al., 2018). However, PHP can be induced in the absence of extracellular Ca^2+^ at the *Drosophila* NMJ (Goel et al., 2017). Moreover, there is some evidence that ion flux through nicotinic acetylcholine receptors is dispensable for PHP at the mouse NMJ (Wang et al., 2018). It is thus currently unclear whether PHP is triggered by reduced Ca^2+^ influx or synaptic function resulting from receptor inhibition.

The molecular mechanisms underlying PHP are understood best at the *Drosophila* NMJ. At this synapse, PHP can be rapidly expressed within minutes after pharmacological glutamate receptor (GluR) inhibition by the GluR antagonist Philanthotoxin-433 (PhTx) (Frank et al., 2006). Similarly, PHP is induced and chronically sustained upon genetic loss of the GluRIIA subunit (Petersen et al., 1997). Electrophysiology-based genetic screens at the *Drosophila* NMJ have implicated several genes in PHP, including the schizophrenia-susceptibility gene *dysbindin* (Dickman and Davis, 2009), or the gene encoding the active zone protein rab3-interacting molecule (RIM) (Müller et al., 2012). Many of the identified genes are required for both, rapid and chronic PHP expression upon pharmacological or genetic receptor impairment, implying that a stereotypic PHP signaling pathway is triggered by acute and chronic receptor perturbation. However, recent work revealed that some genes that promote chronic PHP expression in *GluRIIA* mutants are dispensable for acute PHP expression upon pharmacological receptor impairment (Böhme et al., 2019; James et al., 2019). It was concluded that different PHP phases (rapid vs. chronic) involve different molecular mechanisms. An alternative, yet less explored hypothesis is that PHP is promoted by distinct molecular pathways depending on how the receptors have been perturbed, e.g. genetic ablation vs. pharmacological inhibition, or receptor inhibition with different antagonists.

Here, we tested whether perturbing receptors with different antagonists could activate distinct PHP pathways at the *Drosophila* NMJ. We observed that two GluR antagonists, PhTx and Gyki, induced PHP, whereas the antagonist γDGG did not. Interestingly, PHP induced by PhTx and Gyki exhibited differences in the regulation of active-zone structure, short-term plasticity, and the underlying signaling molecules. While PhTx-induced PHP relied on the known PHP genes *dysbindin* and *RIM*, Gyki-induced PHP was promoted by presynaptic Protein Kinase D (PKD). Moreover, Gyki induced PHP in six mutants previously shown to disrupt PHP upon PhTx treatment. Together, our data suggest that distinct molecular mechanisms mediate PHP in response to GluR inhibition by different antagonists, and that GluR inhibition *per se* is not sufficient for PHP expression

## Results

### GluR perturbation does not induce PHP *per se*

To examine whether different receptor perturbations induce distinct PHP pathways, we investigated rapid PHP at the *Drosophila* larval NMJ after GluR perturbation using three antagonists: PhTx, γ-D-Glutamylglycine (γDGG) and Gyki-53655 (Gyki). Although the mechanisms of antagonism of these three inhibitors have not been examined in *Drosophila*, all of them inhibit *Drosophila* GluRs: While PhTx and γDGG were previously shown to block GluRs at the *Drosophila* NMJ (Frank et al., 2006; Miśkiewicz et al., 2011; Pawlu et al., 2004), Gyki had not been tested on *Drosophila* GluRs. We observed that Gyki treatment reduced the amplitude of miniature excitatory postsynaptic potentials (mEPSPs) in a dose-dependent manner (Fig S1B). Moreover, Gyki decreased the amplitude and accelerated the decay of miniature excitatory postsynaptic currents (mEPSCs; see below in Fig. 4). Interestingly, *Drosophila* GluRs contain several conserved Gyki interaction amino acids identified for rat GluA2 (Fig. S1A). Together, these results are consistent with the idea that Gyki blocks *Drosophila* GluRs.

We next assayed PHP using either PhTx, γDGG or Gyki (see Methods and Fig. S1C). PhTx reduced the median mEPSP amplitude by ~50% (Fig. 1A), indicating robust postsynaptic receptor inhibition. By contrast, there was no significant reduction in the amplitude of action potential (AP)-evoked excitatory postsynaptic potentials (EPSPs) (Fig. 1A, S1D). The disproportionate decrease in mEPSP and EPSP amplitude translates into a significant increase in quantal content (QC = EPSP/mEPSP) (Fig. 1A), suggesting enhanced neurotransmitter release (Frank et al., 2006). The quantal content value of the majority of PhTx-treated NMJs fall within the boundary of the expected quantal content range to restore EPSP amplitudes within ±20% of the median amplitude of untreated EPSPs (Fig. 1B). These data suggest that PhTx-dependent GluR inhibition induces PHP, in line with earlier work (Delvendahl and Müller, 2019).

**Figure 1.**
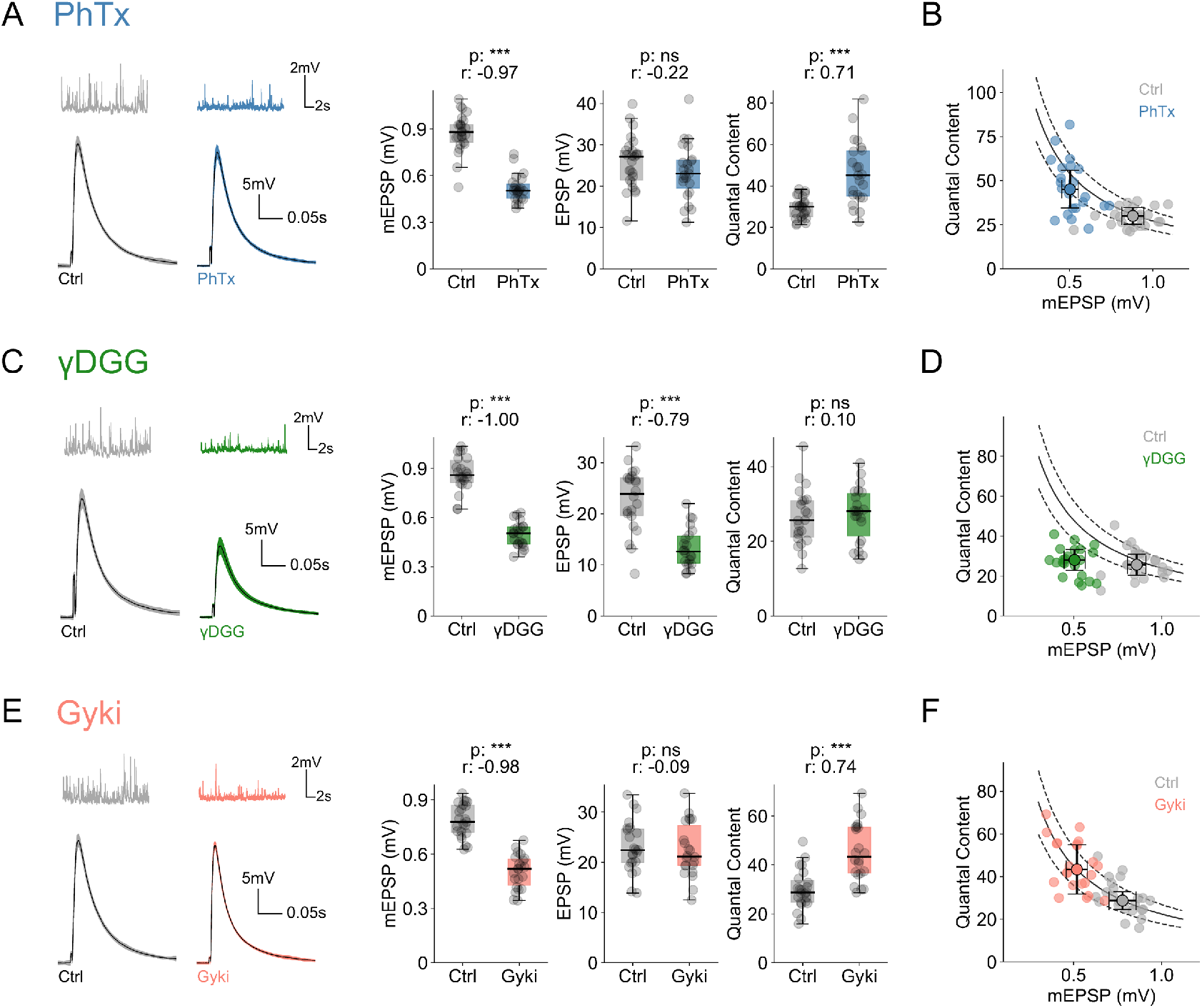
Distinct glutamate receptor perturbations produce different presynaptic homeostatic plasticity (PHP) responses. (A) Representative traces (mEPSPs and EPSPs), mEPSP amplitudes, EPSP amplitudes and quantal content from saline (Ctrl) or PhTx-treated NMJs. n = 25 (Ctrl) vs. 23 (PhTx). (B) Quantal content and median mEPSP amplitude of individual Ctrl and PhTx-treated cells along with group median and median absolute deviation as error bars. (C) Representative traces (mEPSPs and EPSPs), mEPSP amplitudes, EPSP amplitudes and quantal content from saline (Ctrl) or γDGG-treated NMJs. n = 22 (Ctrl) vs. 22 (γDGG). (D) Quantal content and median mEPSP amplitude of individual Ctrl and γDGG-treated cells along with group median and median absolute deviation as error bars. (E) Representative traces (mEPSPs and EPSPs), mEPSP amplitudes, EPSP amplitudes and quantal content from saline (Ctrl) or Gyki-treated NMJs. n = 23 (Ctrl) vs. 22 (Gyki). (F) Quantal content and median mEPSP amplitude of individual Ctrl and Gyki-treated cells along with group median and median absolute deviation as error bars. [Ca2+]e = 0.3 mM. Individual NMJs are shown as gray data points along with the box plot. p-value abbreviated as p and effect size abbreviated as r.

γDGG also reduced the median mEPSP amplitude by ~50% (Fig. 1C). Unlike PhTx, however, EPSP amplitudes dropped proportional to mEPSP amplitudes upon γDGG treatment (Fig. 1C, S1E). Consequently, there was no increase in quantal content (Fig. 1C), and the quantal content value of the majority of γDGG-treated cells did not reach the expected range to restore EPSPs to baseline control amplitudes (Fig. 1D). These observations indicate that the γDGG-induced decrease in mEPSP amplitude is not compensated by PHP.

Treating NMJs with Gyki also significantly decreased mEPSP amplitudes, but not EPSP amplitudes, translating into a significant increase in quantal content (Fig. 1E, S1F). The quantal content value of the majority of Gyki-treated NMJs was close to the expected quantal content to restore baseline EPSPs (Fig. 1F), implying PHP expression upon Gyki treatment. Since PhTx, γDGG and Gyki reduced mEPSP amplitude by a similar magnitude (Fig. 1A,C,E), but only PhTx and Gyki resulted in a homeostatic increase in quantal content, these results indicate that GluR inhibition *per se* is not sufficient to produce PHP.

### Rapid induction and reversal of Gyki-induced PHP

Given that γDGG application did not result in PHP, we focused the rest of our analysis on Gyki and PhTx. Since Gyki has not been used for PHP induction at the *Drosophila* NMJ, we characterized PHP after Gyki treatment.

First, we studied the dynamics of PHP expression in response to Gyki treatment. PhTx induces PHP within 10 minutes after antagonist treatment (Frank et al., 2006). However, little is known about the PHP time course at the *Drosophila* NMJ, because PhTx blocks GluRs in an activity-dependent fashion (Frank et al., 2006). Moreover, PhTx only induces PHP when applied to “semi-intact”, i.e. not fully dissected, larval preparations, which are less amenable to electrophysiological recordings, thus complicating the analysis of PHP dynamics (Frank et al., 2006). In contrast to PhTx, we revealed robust PHP expression upon Gyki treatment of fully-dissected preparations (Fig. S2A), allowing to investigate PHP dynamics.

To probe the dynamics of PHP induction, we quantified mEPSP amplitude, EPSP amplitude and quantal content every 35 seconds before and after Gyki application in the same preparation (Fig. 2A). The temporal resolution of 35 seconds is limited by the time needed to sample mEPSPs for amplitude quantification (see Methods). As expected, mEPSP amplitudes decreased over time after Gyki application (Fig. 2A, 2B, S2B). However, EPSP amplitudes remained largely unchanged (Fig. 2A, 2B, S2B). Consequently, quantal content increased without a measurable delay with regard to mEPSP amplitude reduction (Fig. 2A, 2B, S2B). Considering the temporal resolution of 35 seconds, this implies that EPSP amplitudes are restored continuously as the receptors are being inhibited, pointing towards a low latency of PHP signaling (see Discussion).

**Figure 2.**
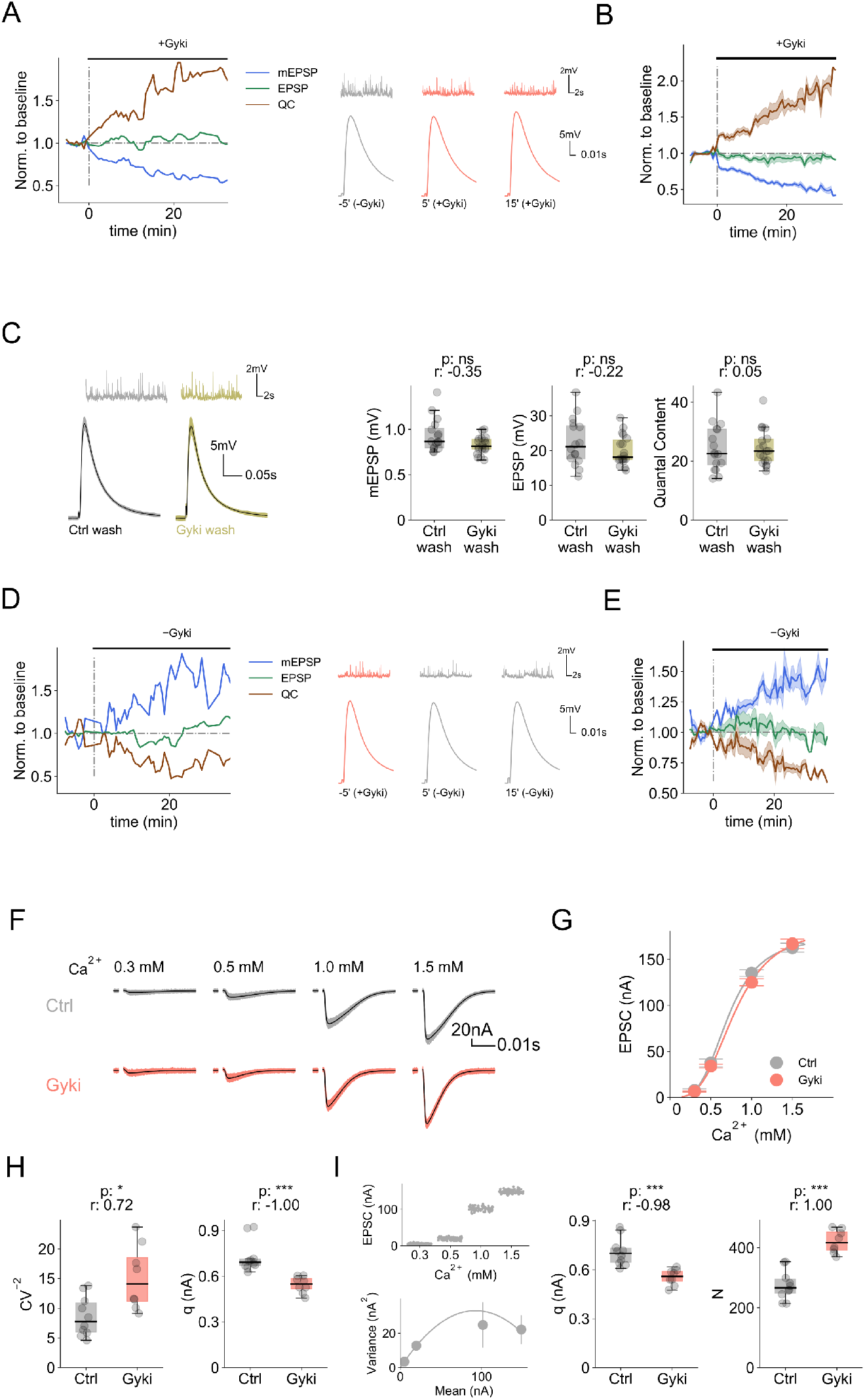
Rapid induction and reversal of Gyki-induced PHP. (A) Left: Normalized median mEPSP amplitude, EPSP amplitude and quantal content of a representative cell as a function of time before and after Gyki treatment, along with. Values are normalized with respective mean baseline values before Gyki application. Right: Representative mEPSP and EPSP traces at specified timepoints. [Ca2+]e = 0.3 mM. (B) Mean normalized mEPSP amplitude, EPSP amplitude and quantal content as a function of time before and after Gyki treatment. The shaded region represents SEM around the mean. Values for individual cells are normalized with the respective mean baseline values before Gyki application. n = 15. (C) Representative traces (mEPSPs and EPSPs), mEPSP amplitudes, EPSP amplitudes and quantal content of NMJs after saline (Ctrl) or Gyki washout. n = 23 (Ctrl wash) vs. 22 (Gyki wash). [Ca2+]e = 0.3 mM. (D) Left: Normalized median mEPSP amplitude, EPSP amplitude and quantal content of a representative cell as a function of time before and after Gyki washout. Values are normalized with respective mean baseline values before Gyki washout. Right: Representative mEPSP and EPSP traces at specified timepoints. [Ca2+]e = 0.3 mM. (E) Mean normalized mEPSP amplitude, EPSP amplitude and quantal content as a function of time before and after Gyki washout. The shaded region represents SEM. Values for individual cells are normalized with the respective mean baseline values before Gyki washout. n = 5. (F) Representative EPSC traces from a saline (Ctrl) and Gyki treated NMJ measured at different extracellular Ca2+ concentrations. (G) EPSC amplitude at different [Ca2+]e for saline-(Ctrl) and Gyki-treated NMJs. n = 16, 16, 51, 27 (Ctrl) vs. 14, 14, 53, 23 (Gyki); Hill’s coefficient = 3.6 (Ctrl) vs. 3.3 (Gyki). (H) Inverse of squared coefficient of variation (CV-2) of EPSC amplitudes recorded at [Ca2+]e = 0.3 mM and estimated quantal size (q) for saline (Ctrl) and Gyki treated NMJs. n = 10 (Ctrl) vs. 8 (Gyki). (I) EPSC amplitude spread as a function of [Ca2+]e, and relation between EPSC variance and mean for a sample variance-mean analysis, along with estimated quantal size (q) and functional release sites (N) for saline-(Ctrl) and Gyki-treated NMJs. n = 10 (Ctrl) vs. 8 (Gyki). Individual NMJs are shown as gray data points along with the box plot. p-value abbreviated as p and effect size abbreviated as r.

We then tested whether Gyki-induced PHP can be reversed upon antagonist washout. NMJs were either incubated with Gyki or saline for 15 minutes, followed by a wash and saline incubation for 15 minutes. mEPSP amplitudes were comparable between Gyki-treated and saline-treated NMJs (Fig. 2C), demonstrating that Gyki reversibly antagonizes *Drosophila* GluRs. In addition, both groups had comparable EPSP amplitudes and quantal content (Fig. 2C), indicating that PHP was reversed upon Gyki removal within 15 minutes. We next explored PHP reversibility dynamics at NMJs that were preincubated with Gyki for 15 minutes. mEPSP, EPSP amplitudes and quantal content were quantified every 35 seconds before and during Gyki washout (Fig. 2D). As expected, mEPSP amplitude increased over time after Gyki washout (Fig. 2D, 2E, S2C). While the evolution of EPSP amplitude over time was relatively variable between cells (Fig. S2C), it remained largely unchanged on average (Fig. 2E). Correspondingly, quantal content decreased without a delay with regard to the increase in mEPSP amplitude given our temporal resolution of 35 seconds (Fig. 2E, S2C), indicating that quantal content was continuously updated as quantal size increased during Gyki washout. Together with the dynamics of PHP induction, these results indicate that PHP at the *Drosophila* NMJ is a highly dynamic and a low-latency process conferring bidirectional synaptic stability on a rapid time scale.

Spontaneous and AP-evoked release could also emerge from different subsets of active zones (Peled et al., 2014). The observation of similar EPSP amplitudes in the absence and presence of Gyki could, in principle, be due to a lack of receptor inhibition at the AP-evoked sites. We therefore next aimed at providing evidence that Gyki also acts on GluRs that are activated by evoked release, and that Gyki indeed potentiates presynaptic release. To this end, we analyzed the amplitudes of AP-evoked excitatory postsynaptic currents (EPSCs) using two-electrode voltage clamp (TEVC) at different extracellular Ca^2+^ concentrations (Fig. 2F). EPSC amplitudes recorded in the absence and presence of Gyki treatment were similar at different Ca^2+^ concentration (Fig. 2F, 2G), suggesting that Gyki does not change the Ca^2+^ sensitivity and Ca^2+^ cooperativity of release, similar to PhTx (Frank et al., 2006). At Gyki-treated NMJs, we observed a significantly increased squared inverse coefficient of variation (CV^−2^) of EPSC amplitudes recorded under conditions of low release probability (*p_r_*) at 0.3 mM Ca^2+^ (Fig. 2H), indicating an increase in quantal content (McLachian, 1978). Furthermore, quantal size (*q*), as estimated as the ratio between mean EPSC amplitude and CV^−2^, was significantly lower for Gyki-treated NMJs, implying postsynaptic receptor inhibition (Fig. 2H). Next, we performed EPSC amplitude variance-mean analysis (Clements and Silver, 2000; Saviane and Silver, 2007) to independently estimate the quantal parameters *q* and functional release site number (*N*) (Fig. 2I; see Methods). Upon Gyki treatment, there was a significant decrease in *q*, and a significant increase *N*, suggesting Gyki-dependent GluR inhibition and presynaptic release potentiation (Fig. 2I). Together, these results independently verify that Gyki treatment impairs GluR function and enhances neurotransmitter release, without major changes in the Ca^2+^ sensitivity or Ca^2+^ cooperativity of release.

### Differential GluR subtype-specificity of Gyki and PhTx

*Drosophila* NMJs harbor two types of GluR complexes, either containing a GluRIIA or a GluRIIB subunit (Marrus et al., 2004). We next tested which receptor complexes are inhibited by Gyki and compared it to PhTx, which is considered to predominantly inhibit GluRIIA-containing receptor complexes (Frank et al., 2006). We recorded mEPSPs from NMJs mutant for either GluRIIA (*GluRIIA^SP16^*; (Petersen et al., 1997)), or GluRIIB (*GluRIIA^SP5^*; (Muttathukunnel et al., 2021)) in the presence and absence of either inhibitor. Whereas mEPSP amplitudes of *GluRIIA* mutant NMJs were largely unaffected by PhTx (Fig. 3A), they significantly decreased in the presence of Gyki (Fig. 3A). mEPSP amplitudes were strongly reduced by both PhTx and Gyki application in *GluRIIB^SP5^* mutants (Fig. 3B). Thus, PhTx predominantly acts on GluRIIA-containing receptors, consistent with previous reports (Frank et al., 2006), whereas Gyki inhibits both, GluRIIA- and GluRIIB-receptor complexes. We conclude that Gyki and PhTx differentially inhibit *Drosophila* GluR types. The differential action on GluR types presents a possibility that Gyki and PhTx may induce PHP via distinct pathways.

**Figure 3.**
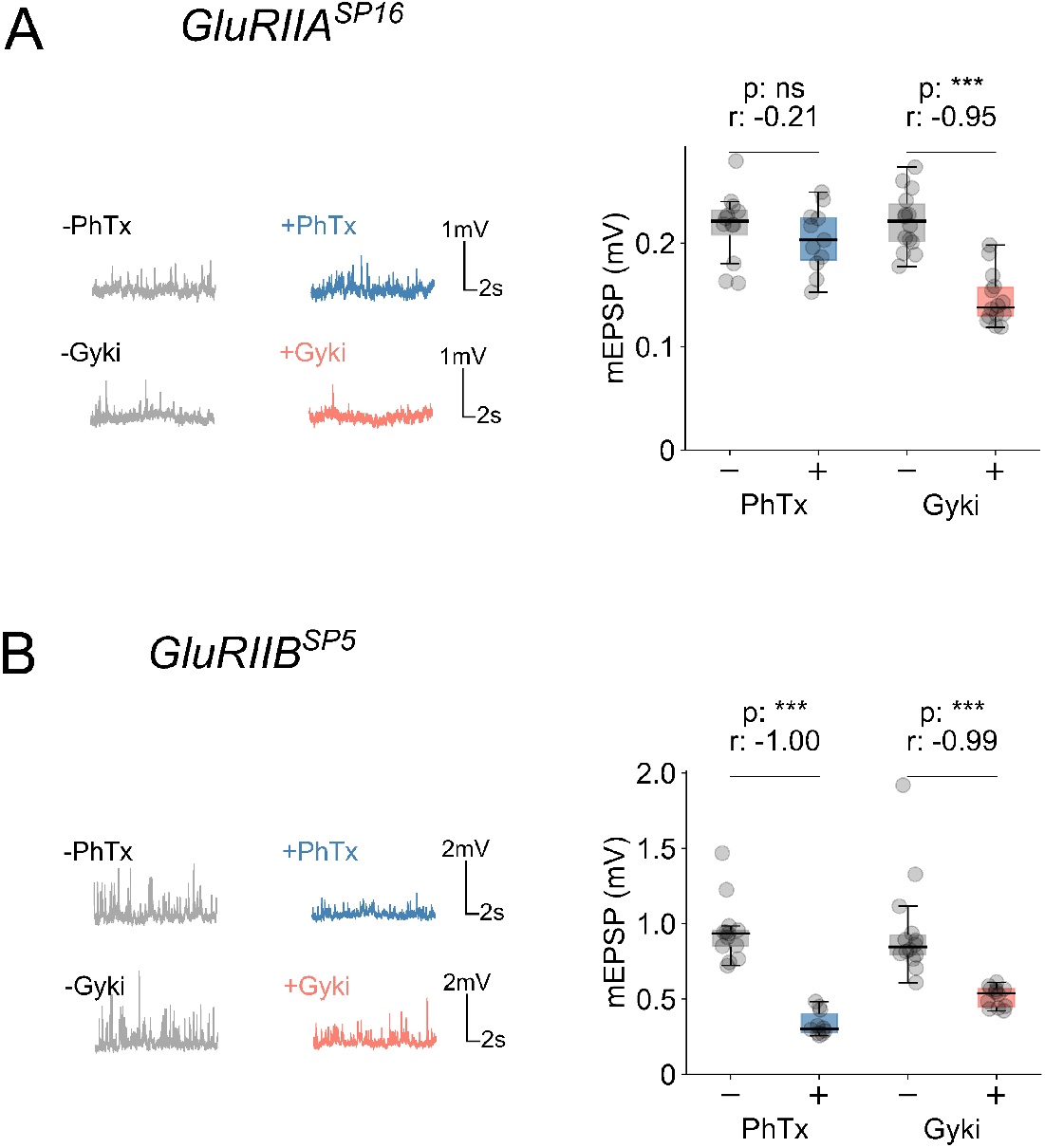
PhTx and Gyki exhibit differential effects on GluRIIA and GluRIIB-containing receptors. (A) Representative mEPSP traces and mEPSP amplitude quantification for PhTx-untreated or treated, and Gyki-untreated or treated GluRIIASP16 mutant NMJs. n = 12 (−PhTx) vs. 11 (+PhTx); n = 14 (−Gyki) vs. 14 (+Gyki). (B) Representative mEPSP traces and mEPSP amplitude quantification for PhTx-untreated or treated, and Gyki-untreated or treated GluRIIBSP5 mutant NMJs. n = 13 (−PhTx) vs. 10 (+PhTx); n = 14 (−Gyki) vs. 12 (+Gyki). Individual NMJs are shown as gray data points along with the box plot. p-value abbreviated as p and effect size abbreviated as r.

### Gyki and PhTx increase RRP size with different effects on short-term plasticity

PhTx induces PHP by increasing readily-releasable pool (RRP) size without apparent effects on short-term dynamics (Ortega et al., 2018; Weyhersmüller et al., 2011). We therefore next did a comparative analysis of these synaptic parameters during PhTx- and Gyki-induced PHP using TEVC (1.0 mM extracellular Ca^2+^). As expected, both, PhTx and Gyki strongly reduced mEPSC amplitudes (Fig. 4A, 4B, left), while EPSC amplitudes remained largely unchanged (Fig. 4A, 4B, middle), resulting in an increase in quantal content (Fig. 4B, right). Moreover, PhTx and Gyki significantly accelerated mEPSC decay kinetics (Fig. 4C, left) and EPSC decay kinetics (Fig. 4C, middle). Interestingly, while Gyki treatment similarly accelerated mEPSC and EPSC decay kinetics (Fig. 4C, right), the PhTx-induced acceleration of mEPSC kinetics was more pronounced than the one of EPSC kinetics (Fig. 4C, right). These data indicate that PhTx, but not Gyki, may potentiate neurotransmitter release during the EPSC decay phase, or that the two antagonists differentially affect GluR properties during evoked transmission (see Discussion).

**Figure 4.**
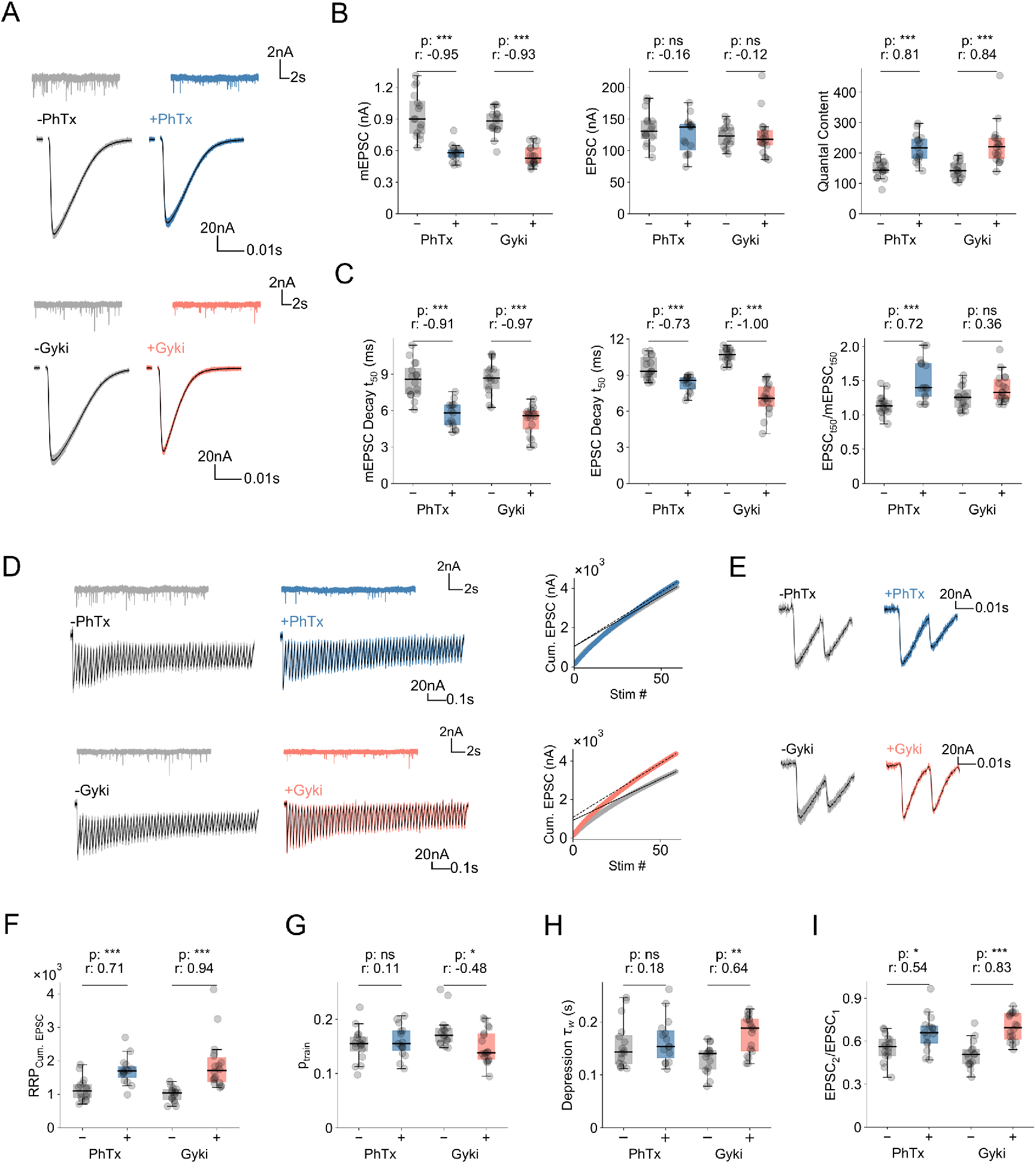
Gyki and PhTx increase RRP size, but with differences in short-term plasticity. (A) Representative traces (mEPSCs and EPSCs), (B) mEPSP amplitudes, EPSC amplitudes, quantal content, (C) mEPSP decay t-half (t50), EPSP decay t-half (t50) and ratio of EPSC t50 and mEPSP t50 for PhTx-untreated or treated, as well as Gyki-untreated or treated NMJs. (D) Representative EPSC trains (60 stimuli at 60 Hz frequency), mEPSC, and linear fitting of last 10 cumulative EPSCs, (E) first two EPSCs of the train from PhTx-untreated or treated, and Gyki-untreated or treated NMJs. (F) RRP size(RRPCum. EPSC) and (G) release probability (ptrain) estimated from cumulative EPSCs of PhTx-untreated or treated, and Gyki-untreated or treated NMJs. (H) Depression time constant of normalized EPSC amplitudes (*τ*w) and (I) Paired-pulse ratio (EPSC2/EPSC1) for PhTx- untreated or treated, and Gyki-untreated or treated NMJs. n = 16 (−PhTx) vs. 15 (+PhTx); n = 15 (−Gyki) vs. 18 (+Gyki). [Ca2+]e = 1.0 mM. Individual NMJs are shown as gray data points along with the box plot. p-value abbreviated as p and effect size abbreviated as r.

We next probed RRP-size modulation during PHP induction by Gyki and compared it to PhTx by back-extrapolating the steady-state of cumulative EPSC amplitudes during repetitive stimulation (60 Hz, 60 stimuli; see Methods; Fig. 4D) (Schneggenburger et al., 1999). RRP size (‘RRP_Cum. EPSC_’) significantly increased after Gyki and PhTx treatment (Fig. 4F). A similar increase in RRP size upon Gyki and PhTx application was obtained with the Elmquist-Quastel method (‘RRP_EQ_’; see Methods; Fig. S3A) (Elmqvist and Quastel, 1965). Hence, Gyki, similar to PhTx, potentiates RRP Size.

Intriguingly, *p_r_*, as assessed by dividing the first EPSC amplitude of a train by the cumulative EPSC amplitude (‘*p_train_*’; (Delvendahl et al., 2019)) was significantly reduced after Gyki, but not PhTx treatment (Fig. 4G, S3B), indicating that Gyki- and PhTx-induced PHP may differentially affect *p_r_*. Consistently, we revealed a significantly slower time course (*τ*_w_; see Methods) of synaptic short-term depression during train stimulation upon Gyki-, but not PhTx treatment (Fig. 4H, S3C), indicative of a *p_r_* decrease after Gyki application. Moreover, Gyki increased the paired-pulse ratio (PPR) of EPSC amplitudes (EPSC_2_/EPSC_1_; 60 Hz; Fig. 4E) compared to untreated controls (Fig. 4E, 4I). The Gyki-induced PPR increase (by 36%) was more pronounced than the PPR change upon PhTx treatment (17%; Fig. 4E, right, 4I). Together, these results show that both, Gyki and PhTx treatment increase RRP size with different effects on *p_r_* and short-term plasticity, indicating the Gyki-induced PHP may involve different mechanisms compared to PhTx-induced PHP (see Discussion).

### PhTx, but not Gyki, upregulates Brp abundance

PhTx application at the *Drosophila* NMJ results in an increase in immunofluorescence intensity of several active zone proteins, including the scaffolding protein bruchpilot (Brp), suggesting increased Brp abundance and/or reorganization (Böhme et al., 2019; Gratz et al., 2019; Weyhersmüller et al., 2011). We therefore examined if Gyki-induced PHP also involves Brp modulation. After PhTx treatment, we detected a significant increase in Brp-fluorescence intensity (Fig. 5B), consistent with previous reports (Böhme et al., 2019; Weyhersmüller et al., 2011). On the other hand, we did not observe an increase in Brp-fluorescence intensity after Gyki treatment (Fig. 5B). Hence, although both PhTx and Gyki inhibit GluRs and induce PHP, only PhTx leads to an increase in Brp abundance, further supporting the idea that different receptor perturbations may trigger different downstream pathways.

**Figure 5.**
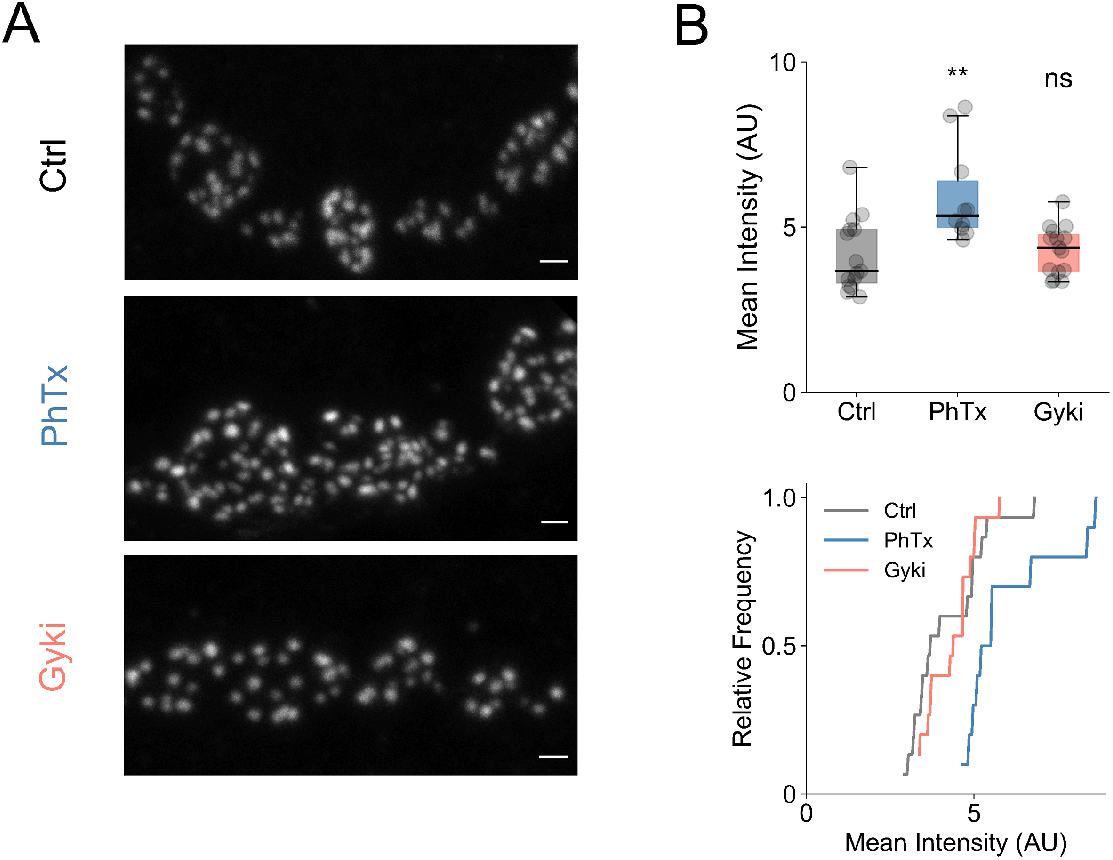
PhTx and Gyki exhibit differences in Brp regulation. (A) Representative confocal images (max. intensity projection) of anti-Brp stained NMJs, treated with saline (Ctrl), PhTx or Gyki. Scale bar: 1 μm. (B) Quantified mean intensities for Ctrl, PhTx or Gyki treated NMJs. Number of NMJs n: 15 (Ctrl), 10 (PhTx), and 15 (Gyki). Displayed statistical significance level obtained from Dunn’s post hoc comparison with Ctrl. Multi-comparison p-value is ~0.003. Individual NMJs are shown as gray data points along with the box plot.

### Gyki induces PHP at mutant NMJs with impaired PhTx-induced PHP

Electrophysiology-based genetic screens in *Drosophila* have identified several genes involved in PhTx-induced PHP (Delvendahl and Müller, 2019). Assaying the role these genes on Gyki-induced PHP could reveal whether PHP induced by Gyki and PhTx involve different molecular mechanisms.

We first tested PHP in a *rim* null mutant, which was shown to have disrupted PhTx-induced PHP (*rim^Δ103^*) (Müller et al., 2012). PhTx application significantly reduced mEPSP and EPSP amplitudes at *rim^Δ103^* mutant NMJs (Fig. 6A, 6B). As the median mEPSP and EPSP amplitudes were reduced by similar fraction upon PhTx treatment in *rim^Δ103^* mutants (Fig. 6C), there was no significant increase in quantal content (Fig. 6B, right, 6D, left), suggesting PHP impairment. Although Gyki treatment significantly decreased mEPSP and EPSP amplitudes in *rim^Δ103^* (Fig. 6A, 6B), the relative decrease in mEPSP amplitude was more pronounced than the relative EPSP amplitude reduction (Fig. 6C). Correspondingly, Gyki significantly enhanced quantal content in *rim^Δ103^* mutants (Fig. 6B, right, 6D, right), with the majority of Gyki-treated NMJs reaching the quantal content range required to restore the median baseline EPSP amplitude (Fig. 6D, right). Thus, while PhTx-induced homeostatic quantal content potentiation requires *rim* (Müller et al., 2012), Gyki enhances quantal content in *rim* mutants, suggesting separable molecular PHP pathways.

**Figure 6.**
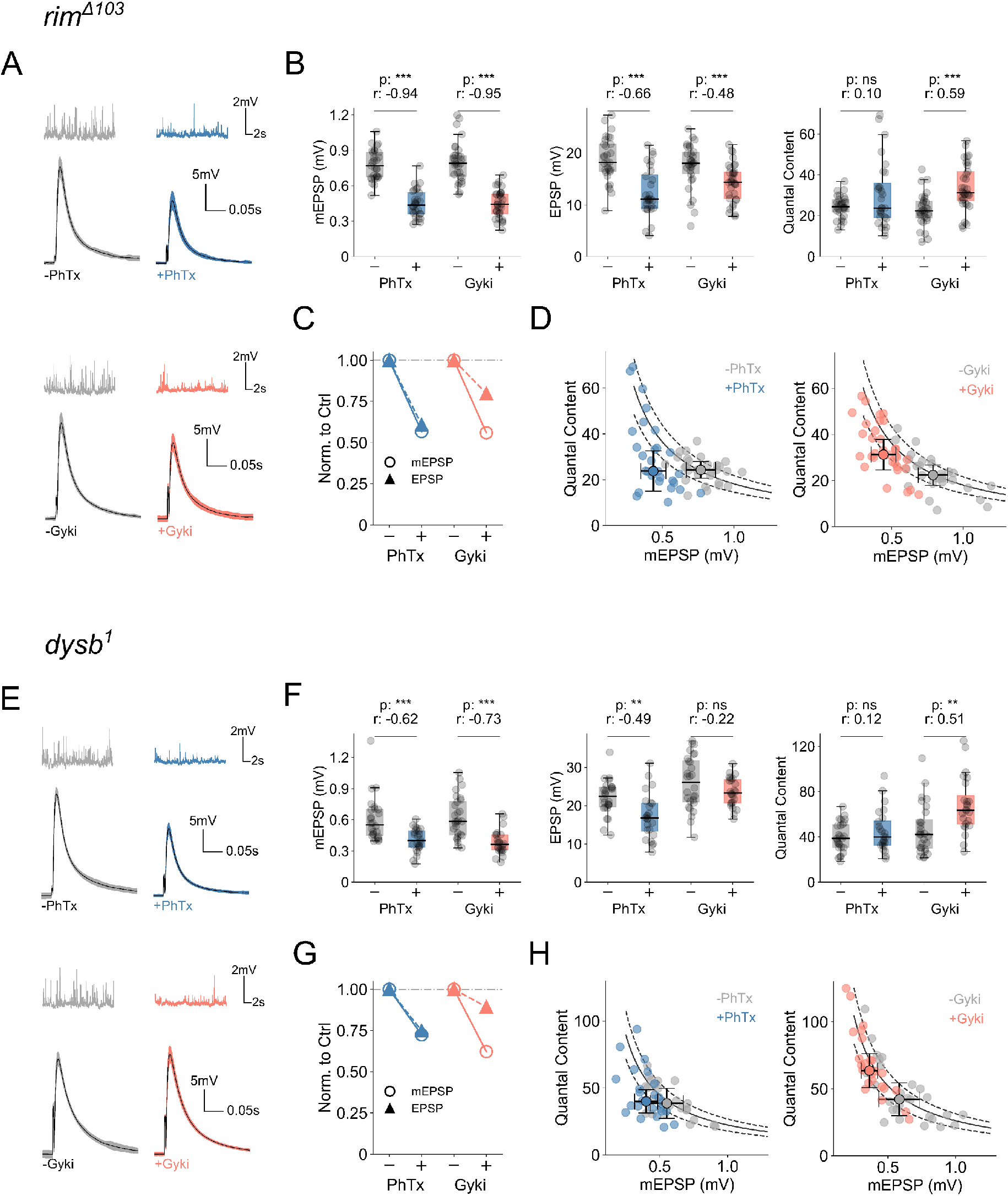
Gyki induces PHP at mutant NMJs with impaired PhTx-induced PHP. (A) Representative traces (mEPSPs and EPSPs), (B) mEPSP amplitudes, EPSP amplitudes, and quantal content for PhTx-untreated or treated, and Gyki-untreated or treated rim mutant NMJs. (C) Comparison of median mEPSP (open circles with solid line) and EPSP (closed circles with dashed line) amplitude without or with either treatment. (D) Quantal content and mEPSP amplitude of individual Ctrl and treated cells along with the median and median absolute deviation as error bars. n = 29 (−PhTx) vs. 28 (+PhTx); n = 30 (−Gyki) vs. 33 (+Gyki). (E) Representative traces (mEPSPs and EPSPs), (F) mEPSP amplitudes, EPSP amplitudes, and quantal content for PhTx-untreated or treated, and Gyki-untreated or treated dysb mutant NMJs. (G) Comparison of median mEPSP (open circles with solid line) and EPSP (closed circles with dashed line) amplitude without or with either treatment. (H) Quantal content and mEPSP amplitude of individual Ctrl and treated cells along with the median and median absolute deviation as error bars. n = 26 (−PhTx) vs. 24 (+PhTx); n = 28 (−Gyki) vs. 24 (+Gyki). [Ca2+]e = 0.4 mM. Individual NMJs are shown as gray data points along with the box plot. p-value abbreviated as p and effect size abbreviated as r.

Next, we probed PHP after loss of *dysbindin* (*dysb*), another gene implicated in PhTx-induced PHP (Dickman and Davis, 2009; Wentzel et al., 2018). Both, PhTx and Gyki treatment significantly reduced mEPSP amplitudes in *dysb^1^* mutants (Fig. 6E, 6F). EPSP amplitudes were significantly decreased upon PhTx application (Fig. 6F), to a similar extent as mEPSP amplitudes (Fig. 6G). On the other hand, EPSP amplitudes were largely unchanged after Gyki treatment (Fig. 6F,G). Consequently, quantal content was unchanged after PhTx application, but significantly increased after Gyki treatment (Fig. 6F, 6H), thereby restoring EPSP amplitudes to baseline levels (Fig. 6H). These data provide genetic evidence that *dysbindin* is required for PhTx-induced PHP, but dispensable for Gyki-induced PHP.

In addition, we assayed Gyki-induced PHP in four additional mutants that were previously shown to disrupt PhTx-induced PHP (Fig. S4). We revealed a significant increase in quantal content upon Gyki application in previously published mutant lines of *DKaiR1D* (Kiragasi et al., 2017), *pickpocket11* (Younger et al., 2013), *multiplexin* (Wang et al., 2014), and *insomniac* (Kikuma et al., 2019), (Fig. S4A-D), indicating intact Gyki-induced PHP in all four cases. Thus, six independent experiments provide evidence that PhTx and Gyki-induced PHP involve distinct molecular mechanisms.

### Presynaptic Protein Kinase D promotes Gyki-induced PHP, but not PhTx-induced PHP

We next carried out a small electrophysiology-based genetic screen to identify molecular players involved in Gyki-induced PHP (Fig. 7A). To this end, we assayed Gyki-induced PHP after genetic perturbation (presynaptic/postsynaptic RNAi expression or validated mutants) of genes encoding for second messenger signaling elements, and previously characterized PHP-related genes. While most transgenic lines displayed a significant increase in quantal content upon Gyki application that restored EPSP amplitudes towards baseline levels (within 20% of the median baseline EPSP; Fig. 7A, solid and dashed lines; see Methods), Gyki treatment did not enhance quantal content in 8 transgenic lines, which represent candidate PHP genes. One of the identified genotypes was an RNAi targeting *Drosophila Protein Kinase D* (*PKD*) presynaptically (*D42-Gal4>UAS-PKD^RNAi^*; Fig. 7A; red circle), a gene not studied in the context of synaptic transmission in *Drosophila* so far. This candidate gene was selected for further analysis.

**Figure 7.**
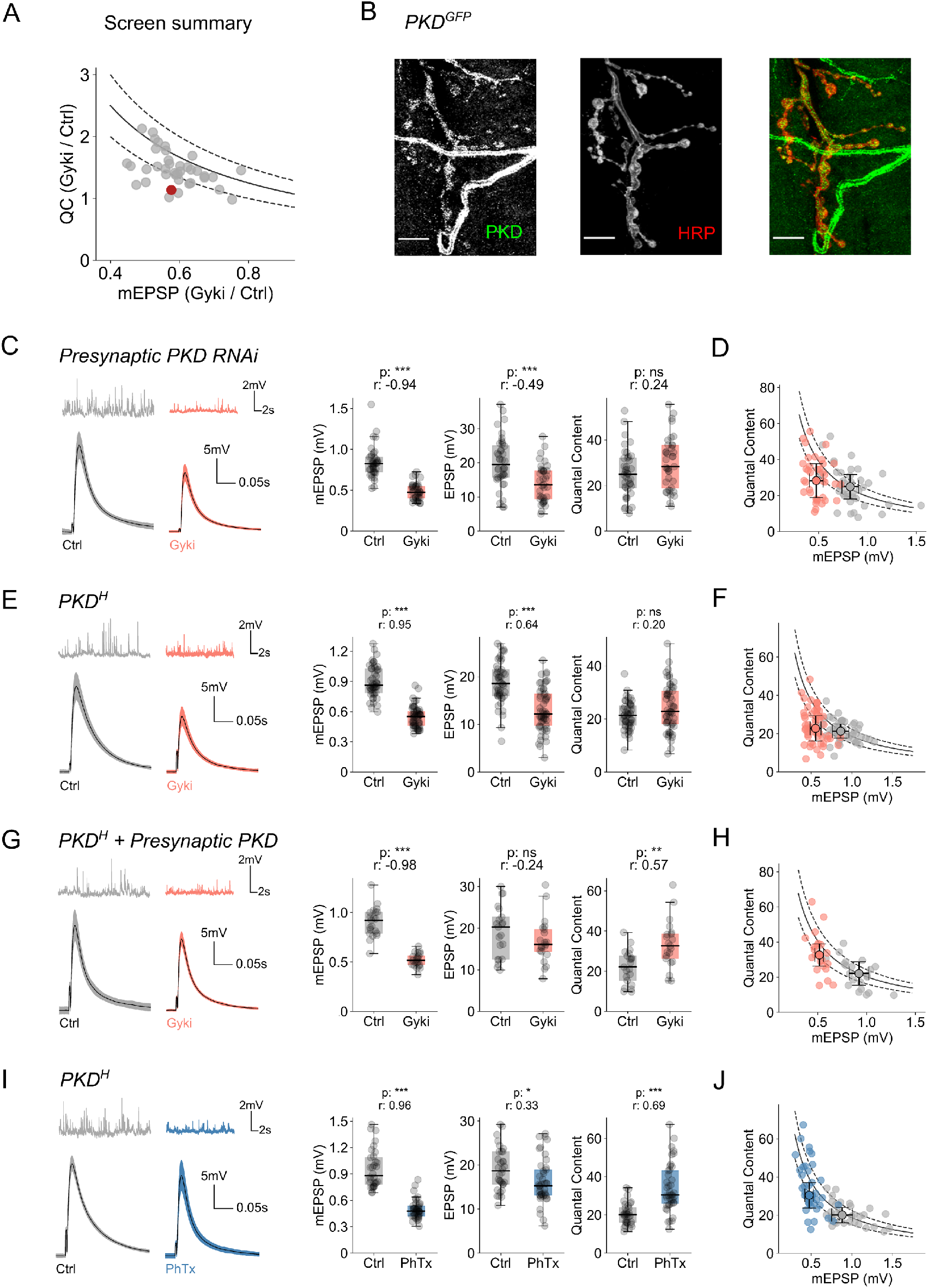
Presynaptic PKD promotes Gyki-induced PHP, but not PhTx-induced PHP. (A) Summary of the screen to identify genes involved in Gyki-induced PHP. Each genotype is represented by a dot representing the relation between ratio of median quantal content (QC) with or without Gyki treatment, and ratio of median mEPSP amplitude with or without Gyki treatment. The solid line, dashed lines below and above the solid line represent functions 1/x, 0.8/x and 1.2/x, respectively. Genotypes falling within the upper and lower dashed lines have similar EPSPs (±20%) with or without Gyki treatment. Genotypes below the lower dashed line have EPSP reduction of more than 20% after Gyki treatment. Red dot is presynaptic PKD knockdown. (B) Expression of fluorescently-tagged endogenous PKD (stained with anti-GFP) and the neuronal membrane marker HRP. PKD fluorescence (green) overlaps with HRP (red) in the merged image. In addition, PKD expression was seen in other non-neuronal compartments, such as trachea. Scale bar = 10 μm. (C) Representative traces (mEPSPs and EPSPs), mEPSP amplitudes, EPSP amplitudes and quantal content for saline (Ctrl) or Gyki-treated NMJs with presynaptic PKD knockdown. (D) Quantal content and mEPSP amplitude of individual Ctrl and treated cells along with the median and median absolute deviation as error bars. n = 40 (Ctrl) vs. 37 (Gyki). (E) Representative traces (mEPSPs and EPSPs), mEPSP amplitudes, EPSP amplitudes and quantal content for saline (Ctrl) or Gyki-treated hypomorphic PKDH mutant NMJs. (F) Quantal content and mEPSP amplitude of individual Ctrl and treated cells along with the median and median absolute deviation as error bars. n = 48 (Ctrl) vs. 55 (Gyki). (G) Representative traces (mEPSPs and EPSPs), mEPSP amplitudes, EPSP amplitudes and quantal content for saline (Ctrl) or Gyki-treated NMJs expressing wild-type PKD presynaptically in the PKDH mutant background. (H) Quantal content and mEPSP amplitude of individual Ctrl and treated cells along with the median and median absolute deviation as error bars. n = 21 (Ctrl) vs. 21 (Gyki). (I) Representative traces (mEPSPs and EPSPs), mEPSP amplitudes, EPSP amplitudes and quantal content for saline (Ctrl) or PhTx-treated hypomorphic PKDH mutant NMJs. (J) Quantal content and mEPSP amplitude of individual Ctrl and treated cells along with the median and median absolute deviation as error bars. n = 32 (Ctrl) vs. 41 (Gyki). [Ca2+]e = 0.3 mM. Individual NMJs are shown as gray data points along with the box plot along with the box plot. p-value abbreviated as p and effect size abbreviated as r.

Although PKD shows a broad expression pattern in *Drosophila* (Maier et al., 2006), its presence at the NMJ had not been investigated. Therefore, we analyzed PKD expression at the larval body wall of third instar larvae using fluorescently-tagged endogenous PKD (*PKD^GFP^*) generated from an intronic MiMIC insertion in the *PKD* gene (Nagarkar-Jaiswal et al., 2015). PKD fluorescence was present in various compartments, including motor neurons, muscles and trachea (Fig. 7B). It was particularly enriched in close proximity to the neural membrane marker horseradish peroxidase (HRP), demonstrating PKD expression in neurons and the synaptic compartments.

We next aimed at verifying a role for *PKD* in Gyki-induced PHP, and at revealing in which synaptic compartment *PKD* acts during PHP using genetic analysis. Gyki application significantly reduced mEPSP and EPSP amplitudes by a similar degree at NMJs expressing *PKD^RNAi^* presynaptically (Fig. 7C). Consequently, there was no significant increase in median quantal content upon Gyki treatment after presynaptic *PKD^RNAi^* expression (Fig. 7C), with the majority of NMJs failing to reach quantal content values required to restore EPSPs to median baseline levels (Fig. 7D). By contrast, Gyki potentiated quantal content in the genetic controls (*UAS-PKD^RNAi^*/+; Fig. S5A), suggesting PHP expression. Furthermore, NMJs expressing *PKD^RNAi^* postsynaptically (*Mef2-Gal4>UAS-PKD^RNAi^*) also exhibited an increase in quantal content after Gyki treatment (Fig. S5B). Together, these results indicate a presynaptic role of PKD in Gyki-induced PHP. Gyki-induced PHP was not blocked after presynaptic *PKD^RNAi^* expression at elevated extracellular Ca^2+^ (Fig. S5C), similar to genes that predominantly promote PhTx-induced PHP at low extracellular Ca^2+^ concentration (Genç et al., 2017; Hauswirth et al., 2018; Kiragasi et al., 2017).

Next, we independently verified a role of PKD in Gyki-induced PHP using a previously characterized hypomorphic allele (*PKD^H^*) (Ashe et al., 2018). Gyki treatment of *PKD^H^* NMJs resulted in a similar fractional reduction in mEPSP and EPSP amplitudes (Fig. 7E), translating into no significant changes in quantal content after Gyki treatment (Fig. 7E, 7F). By contrast, isogenic controls (*yw^MiMIC^*) exhibited a significant increase in quantal content upon Gyki treatment (Fig. S5D). We then tested whether the PHP impairment in *PKD^H^* mutants could be rescued by synaptic compartment-specific expression of a wild-type *PKD* transgene (*UAS-PKD^WT^*) (Maier et al., 2007) in the *PKD^H^* mutant background. Gyki treatment of *PKD^H^* NMJs expressing *PKD* presynaptically (*OK371-Gal4>UAS-PKD^WT^*; *PKD^H^*) resulted in a significant decrease in mEPSP amplitudes (Fig. 7G). However, unlike in the case of *PKD^H^* (Fig. 7E) and the rescue control (*UAS-PKD^WT^*; *PKD^H^* (No Gal4), Fig. S5E), there was no reduction in EPSP amplitude, and thus a significant increase in quantal content, indicating robust PHP expression (Fig. 7G, 7H). By contrast, Gyki application did not result in a significant quantal content increase after muscle-specific *PKD* expression in *PKD^H^* mutants (*Mef2-Gal4>UAS-PKD^WT^*; *PKD^H^*) (Fig. S5F). Thus, presynaptic, but not postsynaptic *PKD* expression restores PHP in the *PKD^H^* mutant background, supporting our conclusion of a presynaptic function of PKD during Gyki-induced PHP.

In principle, the PHP defect in PKD mutants may arise from impaired NMJ development or baseline synaptic transmission. We therefore next investigated possible effects of presynaptic *PKD^RNAi^* expression on NMJ morphology and synaptic transmission (Fig. S6). The total NMJ surface area, as quantified by the neuronal membrane marker HRP, and the number of active zones, as quantified by the number of Brp puncta, were not affected after presynaptic *PKD^RNAi^* expression (Fig. S6A). Moreover, we did not detect apparent differences in miniature and evoked synaptic transmission between presynaptic *PKD^RNAi^*-expressing NMJs and controls (Fig. S6B). Furthermore, RRP size (Fig. S6C), release probability (Fig. S6C), synaptic depression kinetics (Fig. S6D), and paired-pulse ratios were similar between presynaptic *PKD^RNAi^*-expressing NMJs and controls (Fig. S6E). We conclude that presynaptic PKD plays no major role in the regulation of gross NMJ morphology and baseline synaptic transmission, and that the PHP defect is unlikely a secondary consequence of major changes in NMJ development or impaired baseline synaptic transmission.

Finally, we investigated whether PKD is also involved in PhTx-induced PHP by probing synaptic responses of *PKD^H^* mutant NMJs after PhTx application (Fig. 7I). PhTx strongly reduced mEPSP amplitudes in *PKD^H^* mutants with only a small effect on EPSP amplitudes, translating into a significant increase in quantal content (Fig. 7I, 7J). Hence, PhTx treatment in *PKD^H^* mutants results in an increase in quantal content, suggesting that PKD is not required for PhTx-induced PHP expression. This differential requirement of PKD for Gyki- and PhTx-induced PHP further supports the notion that Gyki and PhTx induce PHP via distinct molecular mechanisms.

## Discussion

We here demonstrate that neurotransmitter receptor impairment by different GluR antagonists (PhTx and Gyki) induce PHP via distinct mechanisms, including differential inhibition of GluR subtypes, differential modulation of an active-zone scaffold, as well as short-term plasticity during PHP. Importantly, we provide genetic evidence that separable molecular mechanisms promote PHP in response to Gyki and PhTx treatment. On the other hand, the GluR antagonist γDGG did not produce PHP despite a robust inhibition of postsynaptic receptors. Together, our data suggest that distinct molecular mechanisms mediate PHP in response to GluR inhibition by different antagonists, and that GluR inhibition *per se* is not sufficient for PHP expression.

Our results contrast with prevalent models of homeostatic synaptic plasticity, which rest on the assumption that synaptic activity changes induce homeostatic signaling (Frank et al., 2006; Turrigiano, 2008). In the case of PHP, the magnitude of the homeostatic increase in presynaptic release scales with the decrease in the amplitude of postsynaptic miniature events (Frank et al., 2006). Based on these data, it was proposed that reduced ion flux through the receptors triggers a homeostatic signaling cascade in the postsynaptic cell, which is relayed to the presynaptic compartment where it adjusts release. A prediction that directly follows is that any receptor perturbation with similar effects on the amplitude of synaptic miniature events produces a homeostatic response via similar underlying mechanisms. We observed that two GluR antagonists, PhTx and Gyki, induced PHP, while one antagonist, γDGG, did not. All three perturbations similarly decreased quantal size, indicating a similar reduction of ion flux through the receptors. Although we cannot exclude the possibility that γDGG may have directly blocked PHP expression, our results suggest that ion flux is unlikely the sole signal responsible for PHP induction at the *Drosophila* NMJ. This agrees with recent observations at the *Drosophila* and mouse NMJ indicating that Ca^2+^ flux through the receptor is likely dispensable for PHP induction (Goel et al., 2017; Wang et al., 2018). How could pharmacological receptor perturbation induce PHP independent of ion flux through the receptor? One intriguing possibility is that perturbation-specific conformational changes of the receptor may be involved in PHP signaling (Goel et al., 2017; Wang et al., 2018). Conformational changes of kainate-, AMPA- and NMDA-type GluRs at mammalian synapses are known to signal independent of ion flux in the context of synaptic plasticity, possibly through metabotropic signaling (Dai et al., 2021; Valbuena and Lerma, 2016). It is also known that different antagonists stabilize different conformational states of AMPA receptors (Balannik et al., 2005; Twomey et al., 2018). Thus, it is conceivable that distinct antagonists may trigger different PHP signaling pathways, depending on the different signaling partners that could be recruited by different conformational states. This hypothesis requires further investigation in relation to molecular signaling underlying PHP induction.

Although PhTx and Gyki robustly induced PHP through an increase in RRP size, we observed several differences between PhTx- and Gyki-induced PHP: First, PhTx and Gyki inhibited different GluR subtypes, with PhTx mainly inhibiting GluRIIA-containing receptors, and Gyki blocking both, GluRIIA- and GluRIIB-containing receptors. Therefore, PhTx-dependent receptor inhibition may trigger PHP by affecting a signaling module associated with GluRIIA-containing receptor complexes, whereas Gyki-induced PHP may be triggered by either, or both GluRIIA- and GluRIIB-dependent signaling. An intriguing possibility to be tested in the future is the possible involvement of GluR subunit-specific interactions of several kinases, such as CaMKII (Incontro et al., 2018), in triggering distinct PHP pathways. Second, while Gyki similarly accelerated the decay of mEPSCs and EPSCs, the EPSC decay was slower than the one of mEPSCs upon PhTx application. The mismatch between the mEPSC and EPSC decay kinetics after PhTx treatment may either result from presynaptic changes, such as the recruitment of synaptic vesicles with a low *p_r_* during the EPSC decay phase (Wentzel et al., 2018), or different effects on GluR desensitization, saturation, or diffusion during evoked release (Takahashi et al., 1995). Regardless, as this mismatch was not observed after Gyki treatment, it either reflects a difference in antagonist action or in Gyki- and PhTx-induced PHP. Third, Gyki, but not PhTx, resulted in altered short-term plasticity, indicative of a decrease in *p_r_* or increased GluR desensitization and/or saturation. Fourth, while PhTx increased Brp-fluorescence intensity, indicating elevated Brp abundance (Weyhersmüller et al., 2011), Gyki did not. Although we cannot exclude Brp modulation at a different time point with regard to Gyki application, this observation suggests differential regulation of this core active-zone protein during PhTx- and Gyki-induced PHP. Fifth, we revealed that Gyki application induced PHP in six mutants that were previously shown to block PHP upon PhTx treatment. Sixth, we identified a gene (PKD) that is required for Gyki-induced PHP, but not PhTx-induced PHP. Thus, although we cannot exclude the possibility of partially overlapping molecular mechanisms, these data demonstrate separable molecular pathways between Gyki- and PhTx-induced PHP.

Diverging homeostatic signaling has been observed in the context of acute versus chronic receptor perturbations. At the *Drosophila* NMJ, mutants with intact acute PHP in response to PhTx application displayed impaired chronic PHP in the *GluRIIC* mutant background (James et al., 2019). However, it is difficult to separate whether these differences emerge because of differences in the nature or the timescale of receptor perturbation. Since our experiments focus on acute perturbations, our results support the idea that distinct homeostatic signaling pathways can be triggered depending on the specific way receptors have been perturbed. PhTx, γDGG and Gyki inhibit AMPA receptors via different modes of antagonism. While PhTx is an activity-dependent pore blocker (Jackson et al., 2011), γDGG acts as a low-affinity competitive antagonist (Liu et al., 1999), and Gyki is an allosteric inhibitor (Balannik et al., 2005). PhTx and γDGG inhibit *Drosophila* GluRs with characteristics similar to AMPA receptor inhibition (Frank et al., 2006; Miśkiewicz et al., 2011; Pawlu et al., 2004). Similarly, the conserved Gyki-interacting residues between rat and *Drosophila* GluRs highlight the possibility that Gyki could also act similarly on *Drosophila* and mammalian GluRs (see Fig. S1A). Thus, PhTx, γDGG and Gyki may inhibit *Drosophila* GluRs via different modes of antagonism, which may be involved in triggering distinct homeostatic signaling pathways. Differential homeostatic signaling has also been observed in the context of firing rate homeostasis, where molecular responses depend on whether the protein or the conductance of a sodium channel protein is eliminated (Kulik et al., 2019). Thus, distinct molecular signaling mechanisms that are specific to the nature, rather than the functional effects of a perturbation, may be a general theme in the context of neuronal homeostasis.

We also implicated presynaptic PKD signaling in synaptic homeostasis. Though PKD has been linked to various intracellular processes, such as vesicle sorting (Baron and Malhotra, 2002), endocytosis (Ellwanger and Hausser, 2013), and the regulation of the actin cytoskeleton (Olayioye et al., 2013), its synaptic function is less explored. Recently, PKD was associated with synaptic plasticity (Oueslati Morales et al., 2020). Our data clearly demonstrate that presynaptic PKD is required for Gyki-induced PHP, but not for PhTx-induced PHP, and establishing that separable molecular pathways govern Gyki- and PhTx-induced PHP. However, PKD was required for PHP only at low extracellular Ca^2+^ concentration, similar to a number of genes supporting PhTx-induced PHP (Genç et al., 2017; Hauswirth et al., 2018; Kiragasi et al., 2017). This indicates that PKD signaling may contribute to the robustness of PHP under conditions of low *p_r_*. It worth highlighting that the PHP defect observed upon presynaptic *PKD* knockdown and in *PKD* hypomorphs may not be fully penetrant, likely because of the genetic nature of the perturbation, and/or genetic compensation by other members of the PKC/CaMK family, as previously described (Maier et al., 2019). This genetic compensation is particularly pronounced in *PKD* null mutants (Maier et al., 2019), which precluded the analysis after complete loss of *PKD* function.

Unlike PhTx (Frank et al., 2006), Gyki induced PHP in the fully-dissected NMJ preparation, which is more amenable to electrophysiology and imaging approaches, and thus allowed probing the dynamics of PHP induction. It is currently unclear why PhTx-induced PHP cannot be observed in the full preparation. One hypothesis is that the muscles/synapses may be stretched in the full preparation, which could lead to a disruption of signaling domains relevant for PhTx-induced PHP (Frank et al., 2006). A second possibility could be the potential disruption of neuro-glial signaling that is important for PhTx-induced PHP in the full preparation (Wang et al., 2020). The fact that Gyki results in PHP after full dissection demonstrates that PHP induction is not limited to the “semi-intact” preparation *per se*. It will be interesting to investigate whether different molecular pathways promoting PHP in response to Gyki and PhTx treatment may be differentially affected by the type of preparation. In this regard, Gkyi induced PHP in *Drosophila multiplexin* (*dmp*) mutants, which were shown to disrupt PhTx-dependent PHP, and neuro-glia signaling (Wang et al., 2020).

Gyki application to the fully-dissected preparation revealed very rapid, low-latency PHP induction kinetics, which maintained EPSP amplitudes constant despite Gyki-induced mEPSP amplitude reduction. However, due to sampling interval in our experiments, we cannot rule out any shorter latencies. In addition, we could not estimate exact latency of the signaling due to slower diffusion-limited receptor inhibition observed in our experiments. The reversible GluR block by Gyki also revealed rapid, low-latency PHP reversal after Gyki washout, consistent with previous observations at mouse cerebellar mossy fiber boutons (Delvendahl et al., 2019), or at mouse NMJs (Wang et al., 2016). Our results imply continuous, bidirectional PHP signaling that compensates for receptor impairment within ~35 seconds, similar to observations at the mouse NMJ (Wang et al., 2016). It will be interesting to explore the molecular mechanisms underlying the induction of distinct PHP pathways, and if distinct molecular pathways also rapidly stabilize synaptic efficacy at other synapses.

## Methods

### Fly Stocks

*Drosophila* stocks were raised at 25°C on normal molasses-containing food. The following genotypes were used: *w^1118^* served as wildtype, *Mef2-Gal4* (kind gift from Monica Zwicky’s lab, University of Zurich), *MHC-Gal4* (kind gift from Graeme Davis’ lab, University of California, San Francisco), *D42-Gal4* (Bloomington Drosophila Stock Center - 8816), *OK371-Gal4* (Bloomington Drosophila Stock Center - 26160), *dysb^1^*, *rim^Δ103^*, *GluRIIA^SP16^* (Petersen et al., 1997), *GluRIIB^SP5^* ((Muttathukunnel et al., 2021) kind gift from Dion Dickman’s lab, University of Southern California), *KaiR1D^2^* (*KaiR1D^MB01010^*; Bloomington Drosophila Stock Center - 22962), *ppk11^Mi^* (*ppk11^MB02012^*; Bloomington Drosophila Stock Center - 23781), *dmp^ΔC^* (kind gift from Graeme Davis’ lab, University of California, San Francisco), *inc^kk3^* (kind gift from Dion Dickman’s lab, University of Southern California), *PKD^H^* (*PKD^MI03619^*; Bloomington Drosophila Stock Center - 37604), *PKD^GFP^* (*PKD^MI09308-GFSTF.0^*; Bloomington Drosophila Stock Center - 60273), *UAS-PKD-RNAi* (TRiP.HMC04179; Bloomington Drosophila Stock Center - 55898), *UAS-PKD^WT^* (kind gift from Anette Preiss’ lab, University of Hohenheim), *yw^MiMIC^* (kind gift from Hugo Bellen’s lab, Baylor College of Medicine). Either the *MHC-Gal4* or *Mef2-Gal4* driver line was used for muscle-specific expression and either *D42-Gal4* or *OK371-Gal4* was used for motor-neuron expression.

### Electrophysiology

Wandering third-instar larvae were dissected in HL3 solution (5 mM KCl, 70 mM NaCl, 10 mM Na-HEPES, 5 mM HEPES, 5 mM trehalose, 115 mM sucrose, 10 mM MgCl_2_) with 0.3 mM CaCl_2_ for sharp-electrode membrane voltage recordings (unless stated otherwise), or 1.0 mM CaCl_2_ for two-electrode voltage clamp (TEVC) recordings. The internal organs, including brain and ventral nerve cord, were carefully removed from the body-wall with intact muscle fibers and innervating motor nerves. Sharp-electrode and TEVC recordings were performed on muscle 6 of segments 3 and 4 with sharp glass electrodes (resistance 10-25 MΩ) using an Axoclamp 900A amplifier (Molecular Devices) and a combination of HS 9A x0.1 and HS 9A x10 headstages. For TEVC recordings, muscle cells were clamped to a membrane potential of −65 mV for EPSCs and −100 mV for mEPSCs. For individual NMJs, mEPSP/Cs were recorded prior to EPSP/Cs induced by stimulating the respective hemi-segmental nerve with single APs (3 ms or 0.3 ms stimulus duration for EPSPs and EPSCs, respectively, 0.3 Hz). A total of 50 EPSCs or 30 EPSPs were recorded to obtain the median EPSC or EPSP value, respectively, for each cell. Train responses were obtained from 5 sweeps of 60 EPSCs at 60 Hz. Variance-mean analysis was done with EPSC measurements from individual cells with varying extracellular Ca^2+^ concentration (0.3 mM, 0.5 mM, 1.0 mM and 1.5 or 3.0 mM; 100 EPSCs at 0.3 Hz per Ca^2+^ concentration).

For PHP experiments, semi-intact (dorsally dissected; with internal organs, brain and ventral nerve cord intact) larvae were incubated with GluR antagonists for ~15 minutes. This was followed by removal of internal organs, brain and ventral nerve cord to obtain a fully-stretched preparation for electrophysiological recordings. Recordings for γDGG and Gyki were performed in the presence of the respective inhibitors. For PhTx, we used the previously described protocol in which PhTx was thoroughly washed out with HL3 after the removal of internal organs, brain and ventral nerve cord (Frank et al., 2006). Due to the irreversible binding of PhTx, a significant fraction of receptors remains inhibited after washout. The following inhibitor concentrations were used: PhTx (20 μM; Cat # sc-255421, Santa Cruz Biotechnology), Gyki (10 μM; Cat # 2555, Tocris) and γDGG (5 mM; Cat # 0112, Tocris). Except for experiments in Fig. 1, all experiments with Gyki were performed by directly applying Gyki to the fully stretched larval preparation. For PHP induction and reversibility timecourse, NMJ activity was recorded as successive sweeps (inter-sweep interval = 5 seconds) of 30 seconds duration containing either one or five EPSPs. Gyki was applied or removed by serial exchange of the bath solution.

### Immunohistochemistry

Third-instar larvae were dissected in HL3 and fixed for 10 minutes in 100% ice-cold ethanol followed by rinsing 3 times with PBS. Larvae were washed afterwards 5x 10 minutes with PBS and 0.1% PBT on a rotating wheel. This was followed by 2 hours blocking with 3% normal goat serum in PBT, before incubating the samples with primary antibodies at 4°C overnight. Larvae were washed 5x 10 min with PBT on the following day and then incubated with secondary antibodies in blocking solution for ~2 hours. This was followed by 5x 10 min wash with PBT before mounting on glass slides with ProLong Gold Antifade (Thermo Fisher Scientific). The following primary antibodies were used: anti-Brp (mouse, nc82; 1:100), anti-Discs large (rabbit, DSHB AB_2617529; 1:1000), anti-GFP (mouse, Life Technologies 3E6; 1:1000) and Alexa-677 conjugated anti-HRP (goat, Jackson ImmunoResearch AB_2338967). The following secondary antibodies were used: Alexa-488 conjugated anti-mouse (Life Technologies A11029; 1:100), Alexa-555 conjugated anti-rabbit (Life Technologies A21429; 1:100), and Atto594 conjugated anti-mouse (ATTO-TEC; 1:100). Images were acquired using an upright Leica Stellaris or inverted Leica SP8 laser scanning microscope (University of Zurich Center for Microscopy and Image Analysis) controlled with LASX software suite (Leica Microsystems, Germany). Images were acquired using either 63x or 100x (for Brp intensity quantification) oil immersion objectives (HC PL APO 1.40 NA Oil STED WHITE; Leica Microsystems, Germany). Emitted light was detected with two HyD detectors in photon counting mode (Leica Microsystems, Germany).

### Data Analysis

Electrophysiology data were analyzed using routines written with scientific python libraries, including numpy, scipy, IPython and neo (Garcia et al., 2014). mEPSP/C events were detected using a template-matching algorithm (Clements and Bekkers, 1997). Due to high noise levels of the high-gain current-injecting headstage, the recorded raw trace was filtered using a second-order Savitzky-Golay filter over 5 ms prior to mEPSC detection. Quantal content was calculated as the ratio of the median EPSP/C amplitude and the median mEPSP/C amplitude for a given cell. The expected quantal content for a given mEPSP amplitude to restore EPSP to the untreated control level was estimated as: 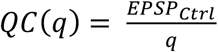, where *q* is the median mEPSP amplitude for a given NMJ, *QC* is quantal content as a function of mEPSP, and *EPSP_Ctrl_* is the median EPSP amplitude of the untreated control group.

As the decay kinetics of NMJ currents do not follow exponential kinetics (Pawlu et al., 2004), mEPSC and EPSC decay kinetics were quantified using the time taken to reach half maximum (t-half) by fitting the decay phase to Hill’s equation: 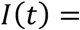 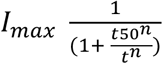, where *I(t)*, *I_max_*, *t*50, and *n* are synaptic current, EPSC amplitude, t-half and steepness coefficient.

For the timecourse experiments, mEPSP amplitude, EPSP amplitude and quantal content were quantified in 35 second bins. The mEPSP, EPSP and quantal content timeseries were then filtered using a first-order Savitzky-Golay filter over three data points.

The readily releasable pool (RRP) size was estimated by the method of back-extrapolation of cumulative EPSC amplitudes measured during stimulus trains (Schneggenburger et al., 1999). Briefly, presynaptic nerve terminals were stimulated with 60 stimuli at 60 Hz and EPSCs were recorded. The cumulative EPSC amplitudes were then plotted as a function of stimulus number. The last 10 data points were fitted with a straight line, and this line was extrapolated to the y-axis intercept. The y-intercept indicates the total response produced by the RRP. RRP size (RRP_Cum. EPSC_) was estimated as the ratio of the y-intercept and the median mEPSC amplitude from the same cell. Release probability (*p_train_*) was estimated by dividing the amplitude of the first EPSC of the train by the total response produced by RRP (y-intercept). RRP size was also independently estimated by analyzing the train responses using the Elmqvist-Quastal method (Elmqvist and Quastel, 1965). Briefly, EPSC amplitudes were plotted as a function of cumulative EPSC amplitudes. The first 5 data points were fitted with a straight line, and this line was extrapolated to the x-axis intercept. The x-axis intercept, in this case, indicates the total response produced by the RRP. RRP size (RRP_EQ_) was estimated as the ratio of the x-intercept and the median mEPSC amplitude from the same cell. Release probability (*p_trainEQ_*) was estimated by dividing the amplitude of the first EPSC of the train by the total response produced by the RRP (x-intercept).

The weighted time constant (*τ*_w_) for synaptic short-term depression was estimated by fitting the normalized EPSC amplitudes during the train with a double exponential function: 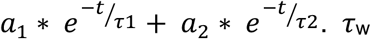 was then estimated using the fitted double exponential function parameters as: 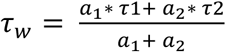. The depression of train EPSC amplitudes at the *Drosophila* NMJs do not follow single exponential decay, and could be better fitted using a double exponential, possibly due to the compound response of Is and Ib synapses, which have different short-term dynamics (Kurdyak et al., 1994).

For variance-mean analysis, EPSC amplitude variance for all Ca^2+^ concentrations were plotted as a function of mean EPSC amplitude for a respective Ca^2+^ concentration and then fitted with: 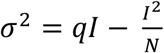, where *σ*, *I*, *q* and *N* are EPSC amplitude variance, mean, quantal size and number of functional release sites, respectively.

For the electrophysiology-based genetic screen to identify candidates involved in Gyki-induced PHP, we scored the relationship between the ratio of quantal content with or without Gyki, and the ratio of mEPSP amplitude with or without Gyki. 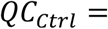 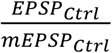 and 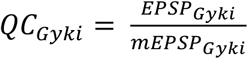, where *QC_Ctrl_*, *EPSP_Ctrl_*, *mEPSP_Ctrl_*, *QC_Ctrl_*, *EPSP_Ctrl_* and *mEPSP_Ctrl_* are median quantal content, median EPSP amplitude and median mEPSP amplitude, without or with Gyki treatment, respectively. Therefore, 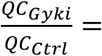 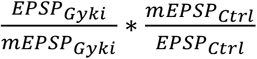. If Gyki treatment results in the restoration of EPSP amplitude within ±20% of the untreated control EPSP level, then 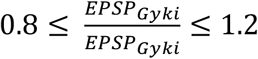. This will follow 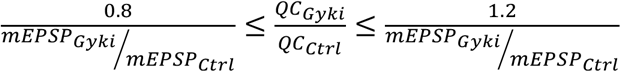. Therefore, we scored 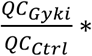 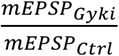 for every genotype, and any genotype with 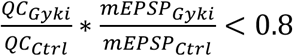 was considered as a candidate gene involved in Gyki-induced PHP.

Microscopy images were analyzed using custom routines written in ImageJ (version 1.51n, National Institutes of Health, USA). Brp fluorescence intensity quantification was performed as follows: First individual Brp puncta were isolated by segmenting binary threshold masks (15% or 35% of the maximum intensity value), background corrected (rolling ball, radius = 1 μm) and filtered (3 x 3 median) maximum intensity projection images. Average intensity values were then calculated for each Brp puncta from raw maximum intensity projection images. HRP-positive regions-of-interest were manually selected and the area were calculated with ImageJ area measurement tool.

### Statistical Analysis

Statistical tests were performed using the stats module of scipy and scikit-posthocs. Two-sided Mann-Whitney U Tests were used to perform statistical comparison, and for multiple data comparison, a Kruskal-Wallis test followed by Dunn’s post hoc comparison was performed. P-values are listed in respective figures as follows; ns : p >= 0.05; * : p < 0.05; ** : p < 0.01; *** : p < 0.001. Effects sizes were quantified as the rank biserial correlation calculated from the U statistics of the Mann-Whitney U Test.

## Acknowledgement

We are grateful to members of the Müller lab for helpful discussions and critical comments. In particular, we would like to thank Igor Delvendahl and Olivier Urwyler for their valuable inputs.

## Funding

This research was funded by a Forschungskredit grant of the University of Zurich and an International Postdoc grant (2018-06813) by the Swedish Research Council to AGN, as well as a Swiss National Science Foundation Assistant Professor grant (PP00P3–15), and an European Research Council Starting grant (SynDegrade-679881) to MM.

## Author Contributions

MM and AGN conceptualized and designed experiments. AGN and PM conducted research and analyzed data. AGN, PM and MM interpreted data. MM and AGN wrote the manuscript.

## Competing Interests

The authors declare no competing interests.

## Data Availability

All data needed to evaluate the conclusions in the paper are present in the paper and the Supplementary Materials.

## Notes

### Competing Interest Statement

The authors have declared no competing interest.

## References

Ashe, S., Malhotra, V., and Raghu, P. (2018). Protein kinase D regulates metabolism and growth by controlling secretion of insulin like peptide. Dev. Biol. 434, 175–185.

Balannik, V., Menniti, F.S., Paternain, A. V, Lerma, J., and Stern-Bach, Y. (2005). Molecular mechanism of AMPA receptor noncompetitive antagonism. Neuron 48, 279–288.

Baron, C.L., and Malhotra, V. (2002). Role of diacylglycerol in PKD recruitment to the TGN and protein transport to the plasma membrane. Science 295, 325–328.

Böhme, M.A., McCarthy, A.W., Grasskamp, A.T., Beuschel, C.B., Goel, P., Jusyte, M., Laber, D., Huang, S., Rey, U., Petzoldt, A.G., et al. (2019). Rapid active zone remodeling consolidates presynaptic potentiation. Nat. Commun. 10, 1–16.

Clements, J.D., and Bekkers, J.M. (1997). Detection of spontaneous synaptic events with an optimally scaled template. Biophys. J. 73, 220–229.

Clements, J.D., and Silver, R.A. (2000). Unveiling synaptic plasticity: a new graphical and analytical approach. Trends Neurosci. 23, 105–113.

Dai, J., Patzke, C., Liakath-Ali, K., Seigneur, E., and Südhof, T.C. (2021). GluD1 is a signal transduction device disguised as an ionotropic receptor. Nature 595.

Delvendahl, I., and Müller, M. (2019). Homeostatic plasticity—a presynaptic perspective. Curr. Opin. Neurobiol. 54, 155–162.

Delvendahl, I., Kita, K., and Müller, M. (2019). Rapid and sustained homeostatic control of presynaptic exocytosis at a central synapse. Proc. Natl. Acad. Sci. U. S. A. 8057, 1–7.

Dickman, D.K., and Davis, G.W. (2009). The schizophrenia susceptibility gene dysbindin controls synaptic homeostasis. Science 326, 1127–1130.

Ellwanger, K., and Hausser, A. (2013). Physiological functions of protein kinase D in vivo. IUBMB Life 65, 98–107.

Elmqvist, D., and Quastel, D.M.J. (1965). A quantitative study of end-plate potentials in isolated human muscle. J. Physiol. 178, 505–529.

Frank, C.A., Kennedy, M.J., Goold, C.P., Marek, K.W., and Davis, G.W. (2006). Mechanisms Underlying the Rapid Induction and Sustained Expression of Synaptic Homeostasis. Neuron 52, 663–677.

Garcia, S., Guarino, D., Jaillet, F., Jennings, T., Pröpper, R., Rautenberg, P.L., Rodgers, C.C., Sobolev, A., Wachtler, T., Yger, P., et al. (2014). Neo: An object model for handling electrophysiology data in multiple formats. Front. Neuroinform. 8, 1–10.

Genç, Ö., Dickman, D.K., Ma, W., Tong, A., Fetter, R.D., and Davis, G.W. (2017). MCTP is an ER-resident calcium sensor that stabilizes synaptic transmission and homeostatic plasticity. Elife 6, e22904.

Genç, Ö., An, J.Y., Fetter, R.D., Kulik, Y., Zunino, G., Sanders, S.J., and Davis, G.W. (2020). Homeostatic plasticity fails at the intersection of autism-gene mutations and a novel class of common genetic modifiers. Elife 9, 1–32.

Goel, P., Li, X., and Dickman, D. (2017). Disparate Postsynaptic Induction Mechanisms Ultimately Converge to Drive the Retrograde Enhancement of Presynaptic Efficacy. Cell Rep. 21, 2339–2347.

Gratz, S.J., Goel, P., Bruckner, J.J., Hernandez, R.X., Khateeb, K., Macleod, G.T., Dickman, D., and O’Connor-Giles, K.M. (2019). Endogenous tagging reveals differential regulation of Ca2+ channels at single active zones during presynaptic homeostatic potentiation and depression. J. Neurosci. 39, 2416–2429.

Haghighi, A.P., McCabe, B.D., Fetter, R.D., Palmer, J.E., Hom, S., and Goodman, C.S. (2003). Retrograde control of synaptic transmission by postsynaptic CaMKII at the Drosophila neuromuscular junction. Neuron 39, 255–267.

Hauswirth, A.G., Ford, K.J., Wang, T., Fetter, R.D., Tong, A., and Davis, G.W. (2018). A postsynaptic PI3K-cII dependent signaling controller for presynaptic homeostatic plasticity. Elife 7, e31535.

Ibata, K., Sun, Q., and Turrigiano, G.G. (2008). Rapid synaptic scaling induced by changes in postsynaptic firing. Neuron 57, 819–826.

Incontro, S., Díaz-Alonso, J., Iafrati, J., Vieira, M., Asensio, C.S., Sohal, V.S., Roche, K.W., Bender, K.J., and Nicoll, R.A. (2018). The CaMKII/NMDA receptor complex controls hippocampal synaptic transmission by kinase-dependent and independent mechanisms. Nat. Commun. 9, 1–21.

Jackson, A.C., Milstein, A.D., Soto, D., Farrant, M., Cull-Candy, S.G., and Nicoll, R.A. (2011). Probing TARP modulation of AMPA receptor conductance with polyamine toxins. J. Neurosci. 31, 7511–7520.

James, T.D., Zwiefelhofer, D.J., and Frank, C.A. (2019). Maintenance of homeostatic plasticity at the drosophila neuromuscular synapse requires continuous IP3-directed signaling. Elife 8, 1–28.

Keck, T., Keller, G.B., Jacobsen, R.I., Eysel, U.T., Bonhoeffer, T., and Hübener, M. (2013). Synaptic scaling and homeostatic plasticity in the mouse visual cortex in vivo. Neuron 80, 327–334.

Kikuma, K., Li, X., Perry, S., Li, Q., Goel, P., Chen, C., Kim, D., Stavropoulos, N., and Dickman, D. (2019). Cul3 and insomniac are required for rapid ubiquitination of postsynaptic targets and retrograde homeostatic signaling. Nat. Commun. 10.

Kiragasi, B., Wondolowski, J., Li, Y., and Dickman, D.K. (2017). A Presynaptic Glutamate Receptor Subunit Confers Robustness to Neurotransmission and Homeostatic Potentiation. Cell Rep. 19, 2694–2706.

Kulik, Y., Jones, R., Moughamian, A.J., Whippen, J., and Davis, G.W. (2019). Dual separable feedback systems govern firing rate homeostasis. Elife 8, 1–27.

Kurdyak, P., Atwood, H.L., Stewart, B.A., and Wu, C. -F (1994). Differential physiology and morphology of motor axons to ventral longitudinal muscles in larval Drosophila. J. Comp. Neurol. 350, 463–472.

Li, X., Goel, P., Chen, C., Angajala, V., Chen, X., and Dickman, D.K. (2018). Synapse-specific and compartmentalized expression of presynaptic homeostatic potentiation. Elife 7, e34338.

Liu, G., Choi, S., and Tsien, R.W. (1999). Variability of neurotransmitter concentration and nonsaturation of postsynaptic AMPA receptors at synapses in hippocampal cultures and slices. Neuron 22, 395–409.

Maier, D., Hausser, A., Nagel, A.C., Link, G., Kugler, S.J., Wech, I., Pfizenmaier, K., and Preiss, A. (2006). Drosophila protein kinase D is broadly expressed and a fraction localizes to the Golgi compartment. Gene Expr. Patterns 6, 849–856.

Maier, D., Nagel, A.C., Gloc, H., Hausser, A., Kugler, S.J., Wech, I., and Preiss, A. (2007). Protein Kinase D regulates several aspects of development in Drosophila melanogaster. BMC Dev. Biol. 7, 1–13.

Maier, D., Nagel, A.C., Kelp, A., and Preiss, A. (2019). Protein Kinase D Is Dispensable for Development and Survival of Drosophila melanogaster. G3 (Bethesda). 9, 2477–2487.

Marrus, S.B., Portman, S.L., Allen, M.J., Moffat, K.G., and DiAntonio, A. (2004). Differential Localization of Glutamate Receptor Subunits at the Drosophila Neuromuscular Junction. J. Neurosci. 24, 1406–1415.

McLachian, E.M. (1978). The statistics of transmitter release at chemical synapses. Int. Rev. Physiol. 17, 49–117.

Miśkiewicz, K., Jose, L.E., Bento-Abreu, A., Fislage, M., Taes, I., Kasprowicz, J., Swerts, J., Sigrist, S., Versées, W., Robberecht, W., et al. (2011). ELP3 controls active zone morphology by acetylating the ELKS family member bruchpilot. Neuron 72, 776–788.

Müller, M., Liu, K.S.Y., Sigrist, S.J., and Davis, G.W. (2012). RIM Controls Homeostatic Plasticity through Modulation of the Readily-Releasable Vesicle Pool. J. Neurosci. 32, 16574–16585.

Muttathukunnel, P., Frei, P., Perry, S., Dickman, D., and Müller, M. (2021). Rapid homeostatic modulation of transsynaptic nanocolumn rings. BioRxiv.

Nagarkar-Jaiswal, S., Lee, P.T., Campbell, M.E., Chen, K., Anguiano-Zarate, S., Gutierrez, M.C., Busby, T., Lin, W.W., He, Y., Schulze, K.L., et al. (2015). A library of MiMICs allows tagging of genes and reversible, spatial and temporal knockdown of proteins in Drosophila. Elife 2015, 1–28.

Olayioye, M.A., Barisic, S., and Hausser, A. (2013). Multi-level control of actin dynamics by protein kinase D. Cell. Signal. 25, 1739–1747.

Ortega, J.M., Genç, Ö., and Davis, G.W. (2018). Molecular mechanisms that stabilize short term synaptic plasticity during presynaptic homeostatic plasticity. Elife 7, 1–28.

Ouanounou, G., Baux, G., and Bal, T. (2016). A novel synaptic plasticity rule explains homeostasis of neuromuscular transmission. Elife 5, 1–20.

Oueslati Morales, C., Ignácz, A., Bencsik, N., Rátkai, A.E., Lieb, W., Eisler, S., Schlett, K., and Hausser, A. (2020). PKD promotes activity-dependent AMPA receptor endocytosis in hippocampal neurons. BioRxiv.

Pawlu, C., DiAntonio, A., and Heckmann, M. (2004). Postfusional control of quantal current shape. Neuron 42, 607–618.

Peled, E.S., Newman, Z.L., and Isacoff, E.Y. (2014). Evoked and spontaneous transmission favored by distinct sets of synapses. Curr. Biol. 24, 484–493.

Perry, S., Han, Y., Das, A., and Dickman, D. (2017). Homeostatic plasticity can be induced and expressed to restore synaptic strength at neuromuscular junctions undergoing ALS-related degeneration. Hum. Mol. Genet. 26, 4153–4167.

Petersen, S.A., Fetter, R.D., Noordermeer, J.N., Goodman, C.S., and DiAntonio, A. (1997). Genetic analysis of glutamate receptors in Drosophila reveals a retrograde signal regulating presynaptic transmitter release. Neuron 19, 1237–1248.

Saviane, C., and Silver, R.A. (2007). Estimation of quantal parameters with multiple-probability fluctuation analysis. Methods Mol. Biol. 403, 303–317.

Schneggenburger, R., Meyer, A.C., and Neher, E. (1999). Released fraction and total size of a pool of immediately available transmitter quanta at a calyx synapse. Neuron 23, 399–409.

Takahashi, M., Kovalchuk, Y., and Attwell, D. (1995). Pre- and postsynaptic determinants of EPSC waveform at cerebellar climbing fiber and parallel fiber to Purkinje cell synapses. J. Neurosci. 15, 5693–5702.

Tatavarty, V., Torrado Pacheco, A., Groves Kuhnle, C., Lin, H., Koundinya, P., Miska, N.J., Hengen, K.B., Wagner, F.F., Van Hooser, S.D., and Turrigiano, G.G. (2020). Autism-Associated Shank3 Is Essential for Homeostatic Compensation in Rodent V1. Neuron 106, 769–777.

Teichert, M., Liebmann, L., Hübner, C.A., and Bolz, J. (2017). Homeostatic plasticity and synaptic scaling in the adult mouse auditory cortex. Sci. Rep. 7, 1–14.

Turrigiano, G.G. (2008). The Self-Tuning Neuron: Synaptic Scaling of Excitatory Synapses. Cell 135, 422–435.

Turrigiano, G.G., Leslie, K.R., Desai, N.S., Rutherford, L.C., and Nelson, S.B. (1998). Activity-dependent scaling of quantal amplitude in neocortical neurons. Nature 391, 892–896.

Twomey, E.C., Yelshanskaya, M. V, Vassilevski, A.A., and Sobolevsky, A.I. (2018). Mechanisms of Channel Block in Calcium-Permeable AMPA Receptors. Neuron 99, 956–968.

Valbuena, S., and Lerma, J. (2016). Non-canonical Signaling, the Hidden Life of Ligand-Gated Ion Channels. Neuron 92, 316–329.

Wang, T., Hauswirth, A.G., Tong, A., Dickman, D.K., and Davis, G.W. (2014). Endostatin Is a Trans-Synaptic Signal for Homeostatic Synaptic Plasticity. Neuron 83, 616–629.

Wang, T., Morency, D.T., Harris, N., and Davis, G.W. (2020). Epigenetic Signaling in Glia Controls Presynaptic Homeostatic Plasticity. Neuron 105, 491–505.e3.

Wang, X., Pinter, M.J., and Rich, M.M. (2016). Reversible Recruitment of a Homeostatic Reserve Pool of Synaptic Vesicles Underlies Rapid Homeostatic Plasticity of Quantal Content. J. Neurosci. 36, 828–836.

Wang, X., McIntosh, J.M., and Rich, M.M. (2018). Muscle Nicotinic Acetylcholine Receptors May Mediate Trans-Synaptic Signaling at the Mouse Neuromuscular Junction. J. Neurosci. 38, 1725–1736.

Wentzel, C., Delvendahl, I., Sydlik, S., Georgiev, O., and Müller, M. (2018). Dysbindin links presynaptic proteasome function to homeostatic recruitment of low release probability vesicles. Nat. Commun. 9, 267.

Weyhersmüller, A., Hallermann, S., Wagner, N., and Eilers, J. (2011). Rapid Active Zone Remodeling during Synaptic Plasticity. J. Neurosci. 31, 6041–6052.

Younger, M.A., Müller, M., Tong, A., Pym, E.C., and Davis, G.W. (2013). A presynaptic ENaC channel drives homeostatic plasticity. Neuron 79, 1183–1196.

