## Supplementary Figures for "Distinct molecular pathways govern presynaptic homeostatic plasticity"

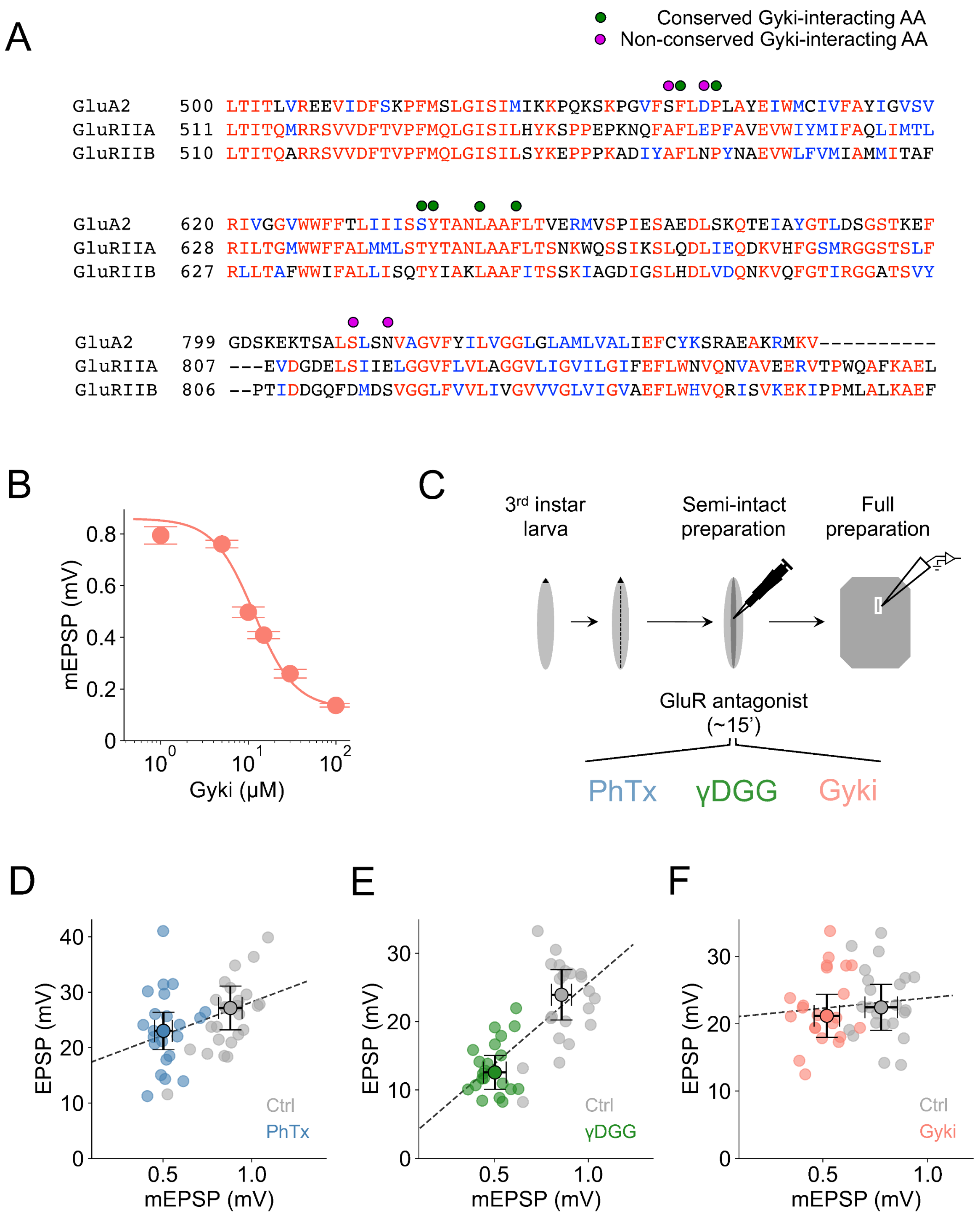

**Fig. S1**. Gyki inhibits *Drosophila* GluRs and induces PHP. (**A**) Multiple sequence alignment of rat GluA2 subunit with *Drosophila GluRIIA* and *GluRIIB* generated from Clustal Omega at EBI web interface using default parameters. Green dots mark columns with conserved Gyki-interacting amino acids (AA), and purple dots mark columns with non-conserved Gyki-interacting residues. Alignment fragments containing Gyki-interacting amino acids (AA) are displayed. (**B**) Dose response curve for Gyki-dependent reduction in mEPSP amplitude. Data are displayed as mean±SEM, whereas the line represents a Hill's fit. $IC_{50}$ = 11.6 μM and Hill's coefficient = 1.8. n = 7, 3, 23, 17, 10, 8 for the respective increasing concentrations. (**C**) Schematic of experimental design for PHP induction. (**D**) EPSP amplitude vs. mEPSP amplitude for individual cells without or with PhTx, (**E**) γDGG, and (**F**) Gyki treatment. The dotted line represents a linear regression fit. Dots with error bars represent median and median absolute deviation of the respective group. Note the increased slope after γDGG treatment. n same as in Fig. 1.

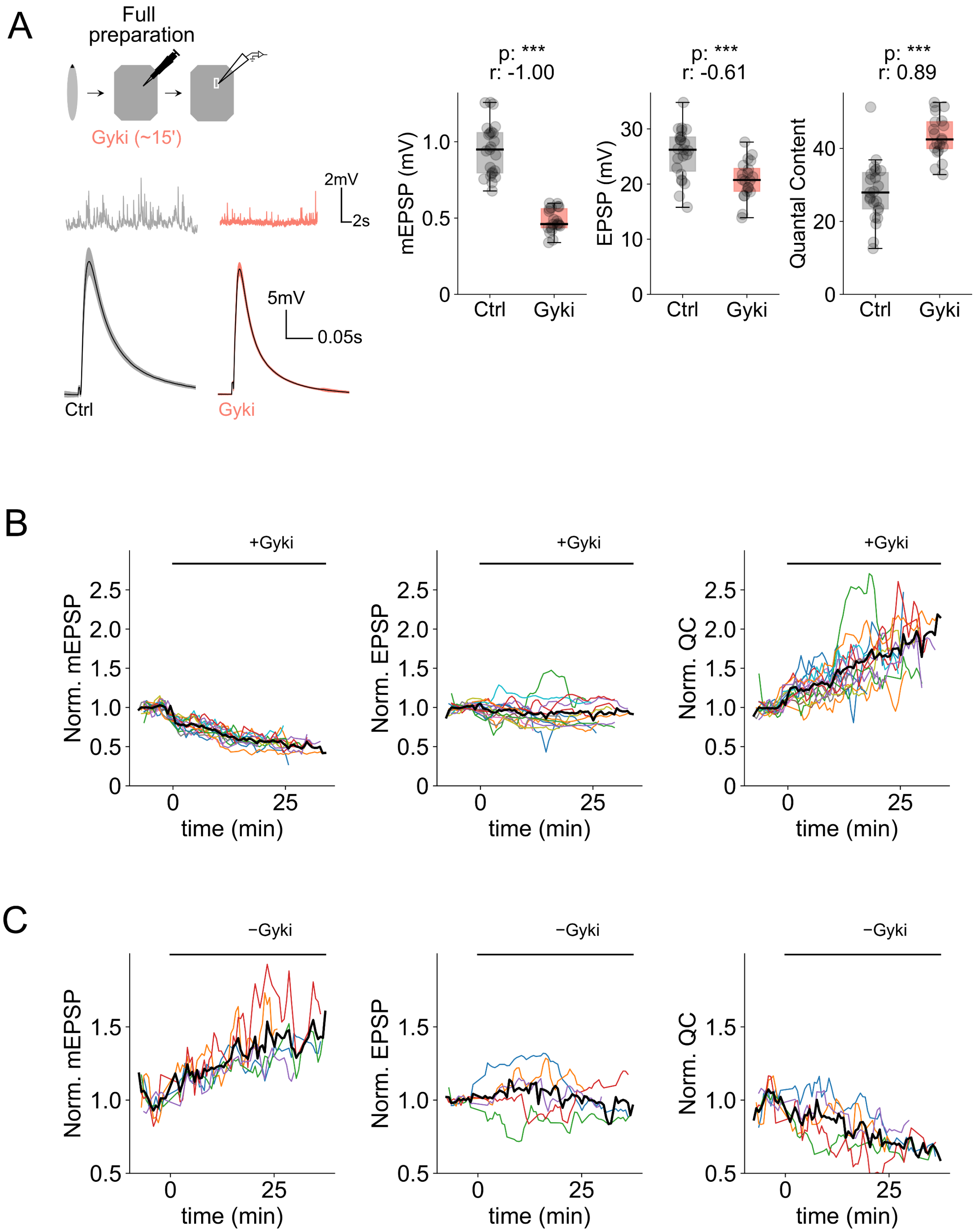
A
Full preparation
Gyki (~15')
2mV
2s
5mV
0.05s
Ctrl
Gyki
p: ***
r: -1.00
mEPSP (mV)
p: ***
r: -0.61
EPSP (mV)
p: ***
r: 0.89
Quantal Content
B
+Gyki
Norm. mEPSP
Norm. EPSP
Norm. QC
time (min)
C
−Gyki
Norm. mEPSP
Norm. EPSP
Norm. QC
time (min)

**Fig. S2**. Further characterization of Gyki. (**A**) Schematic of Gyki treatment on fully-dissected larval preparation along with representative traces (mEPSPs and EPSPs), mEPSP amplitudes, EPSP amplitudes, quantal content from saline (Ctrl) or Gyki-treated NMJs. n = 30 (Ctrl) vs. 32 (Gyki). Individual NMJs are shown as gray data points along with the box plot. (**B**) Normalized mEPSP amplitude, EPSP amplitude and quantal content of individual cells as a function of time before and after Gyki treatment. Values for individual cells are normalized with the respective mean baseline values before Gyki application. Thick black traces are mean of all cells. n = 15; thick black traces: mean. (**C**) Normalized mEPSP amplitude, EPSP amplitude and quantal content of individual cells as a function of time before and after Gyki washout. Values for individual cells are normalized with the respective mean baseline values before Gyki washout. n = 5; thick black traces: mean. $[Ca^{2+}]_e$ = 0.3 mM. p-value abbreviated as p and effect size abbreviated as r.

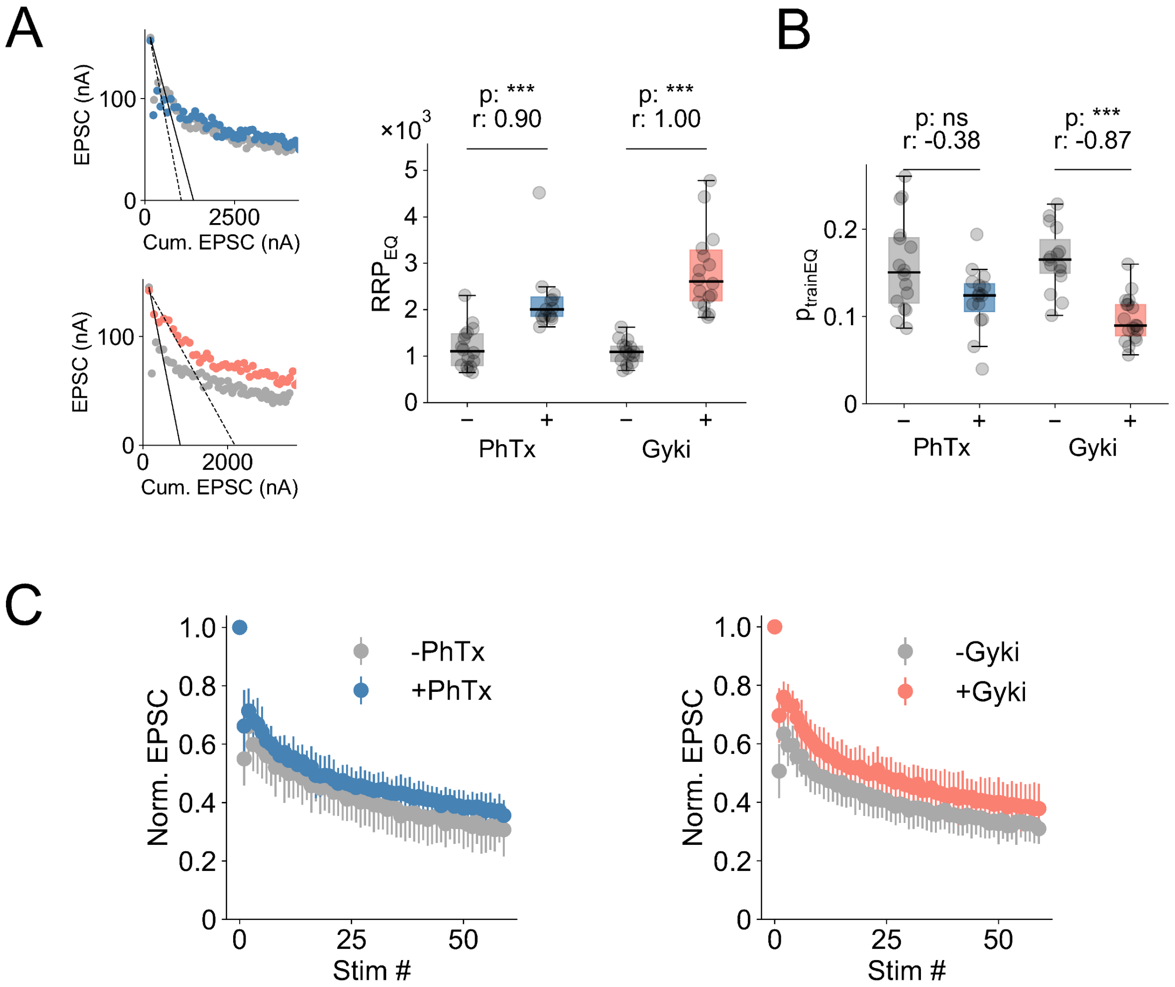


**Fig. S3**. Gyki and PhTx increase RRP size, but with differences in short-term plasticity. (**A**) Representative linear fitting of first 5 EPSCs as a function of cumulative EPSCs to estimate RRP size with the EQ method, along with the estimated RRP size($RRP_{EQ}$) and (**B**) release probability ($p_{trainEQ}$) for PhTx-untreated or treated, and Gyki-untreated or treated NMJs. Individual NMJs are shown as gray data points along with the box plot. (**C**) Normalized mean EPSC amplitudes of all NMJs for PhTx-untreated or treated, and Gyki-untreated or treated experimental groups. Error bars represent standard deviation. n = 16 (-PhTx) vs. 15 (+PhTx); n = 15 (-Gyki) vs. 18 (+Gyki). $[Ca^{2+}]_e$ = 1.0 mM. p-value abbreviated as p and effect size abbreviated as r.

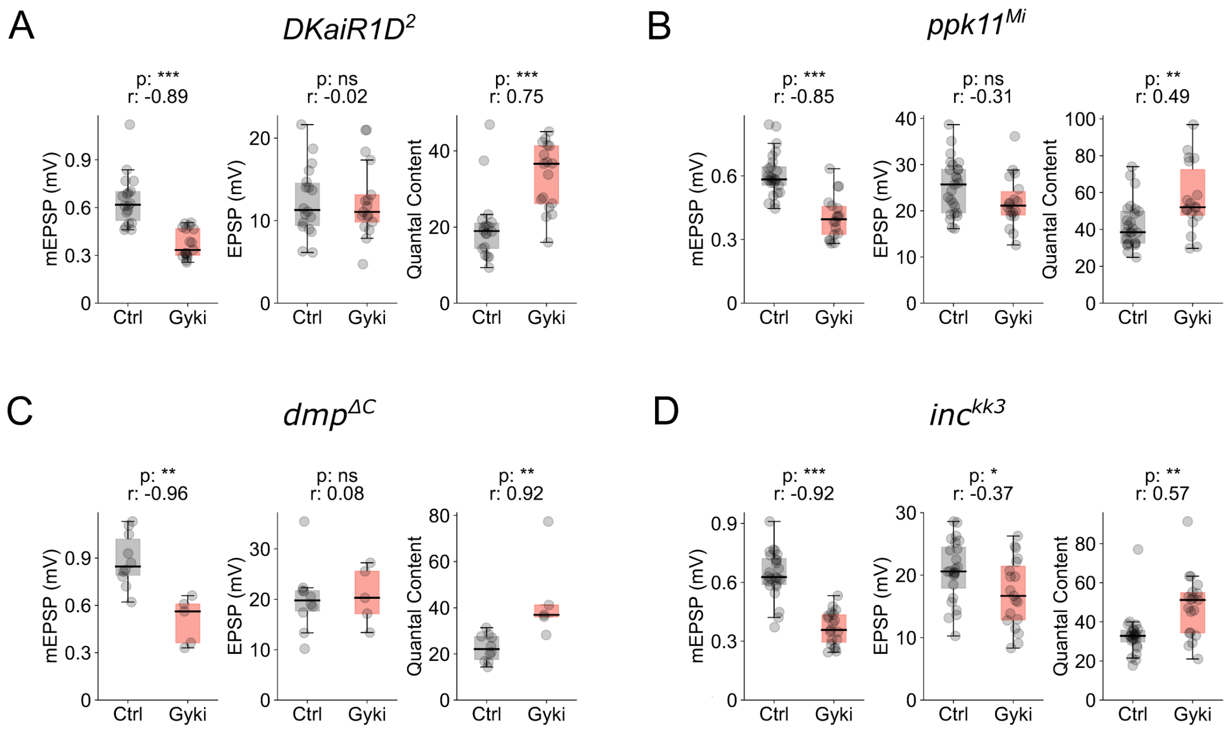


**Fig. S4**. Gyki induces PHP at NMJs mutant for PHP related genes characterized for PhTx. (**A**) Representative traces (mEPSPs and EPSPs), mEPSP amplitudes, EPSP amplitudes, and quantal content of saline (Ctrl) or Gyki-treated *kainate receptor subunit R1D, DKaiR1D*, mutant NMJs. n = 18 (Ctrl) vs. 17 (Gyki). (**B**) Representative traces (mEPSPs and EPSPs), mEPSP amplitudes, EPSP amplitudes, and quantal content of saline (Ctrl) or Gyki-treated *pickpocket11, ppk11*, mutant NMJs. n = 25 (Ctrl) vs. 18 (Gyki). (**C**) Representative traces (mEPSPs and EPSPs), mEPSP amplitudes, EPSP amplitudes, and quantal content of saline (Ctrl) or Gyki-treated *multiplexin, dmp*, mutant NMJs. n = 10 (Ctrl) vs. 5 (Gyki). (**D**) Representative traces (mEPSPs and EPSPs), mEPSP amplitudes, EPSP amplitudes, and quantal content of saline (Ctrl) or Gyki-treated *insomniac, inc*, mutant NMJs. n = 24 (Ctrl) vs. 19 (Gyki). $[Ca^{2+}]_e$ = 0.3 mM. Individual NMJs are shown as gray data points along with the box plot. p-value abbreviated as p and effect size abbreviated as r.

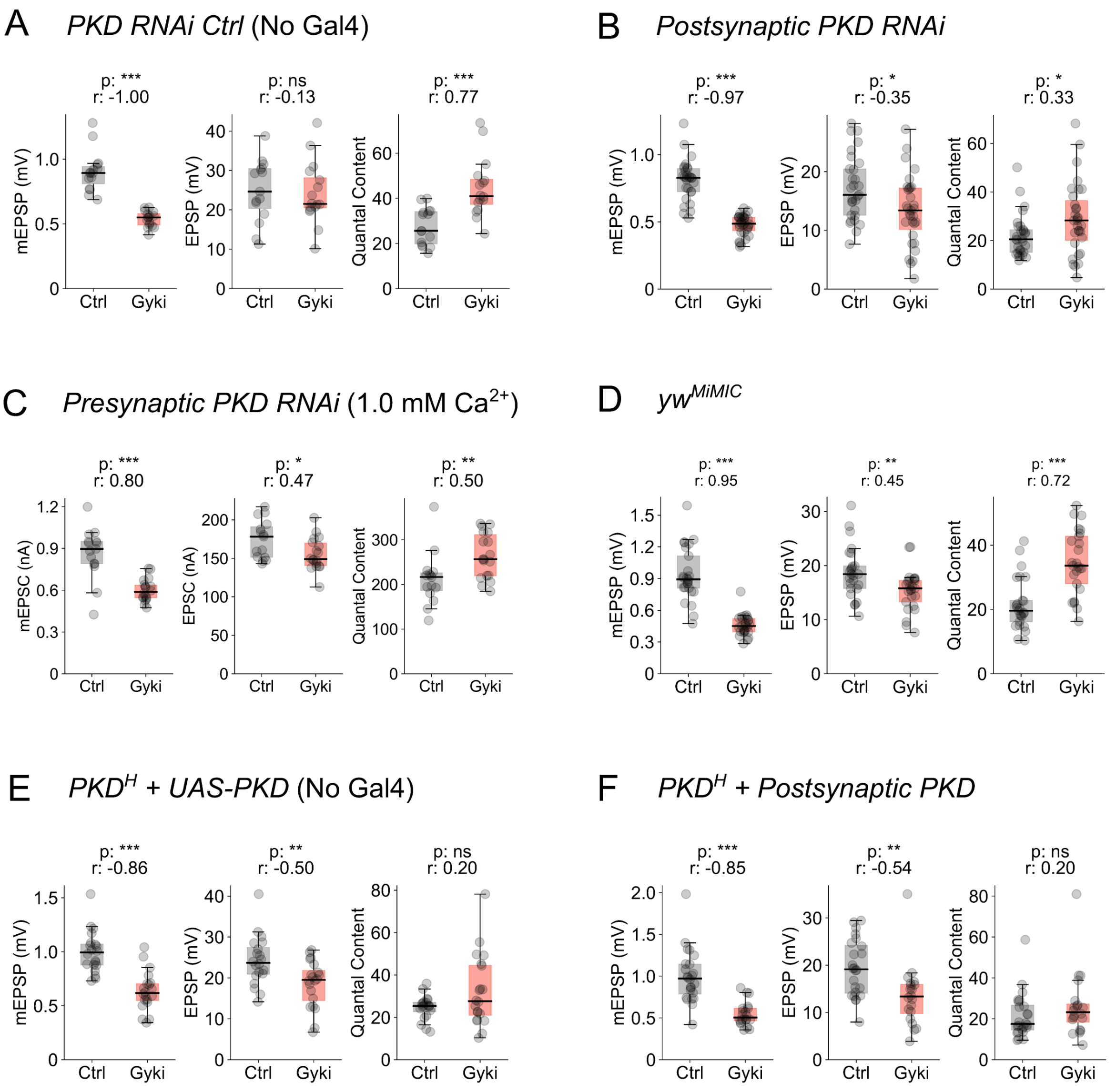


**Fig. S5**. Genetic controls for the role of presynaptic PKD in Gyki-induced PHP. (**A**) mEPSP amplitudes, EPSP amplitudes and quantal content for saline (Ctrl) or Gyki-treated NMJs from *UAS-PKD*$^{RNAi}$ fly line without any Gal4. n = 15 (Ctrl) vs. 16 (Gyki). (**B**) mEPSP amplitudes, EPSP amplitudes and quantal content for saline (Ctrl) or Gyki-treated NMJs with postsynaptic PKD knockdown. n = 29 (Ctrl) vs. 31 (Gyki). (**C**) mEPSP amplitudes, EPSP amplitudes and quantal content for saline (Ctrl) or Gyki-treated NMJs with presynaptic PKD knockdown at $[Ca^{2+}]_e$ = 1.0 mM. n = 15 (Ctrl) vs. 18 (Gyki). (**D**)

mEPSP amplitudes, EPSP amplitudes and quantal content for saline (Ctrl) or Gyki-treated isogenic MiMIC control ($yw^{MiMIC}$) NMJs for $PKD^{H}$. n = 25 (Ctrl) vs. 25 (Gyki). (**E**) mEPSP amplitudes, EPSP amplitudes and quantal content for saline (Ctrl) or Gyki-treated NMJs from larva containing *UAS-PKD* in the $PKD^{H}$ mutant background without any Gal4. n = 20 (Ctrl) vs. 20 (Gyki). (**F**) mEPSP amplitudes, EPSP amplitudes and quantal content for saline (Ctrl) or Gyki-treated NMJs expressing wild-type PKD postsynaptically in the $PKD^{H}$ mutant background. n = 24 (Ctrl) vs. 20 (Gyki). $[Ca^{2+}]_e$ = 0.3 mM, unless stated otherwise. Individual NMJs are shown as gray data points along with the box plot. p-value abbreviated as p and effect size abbreviated as r.

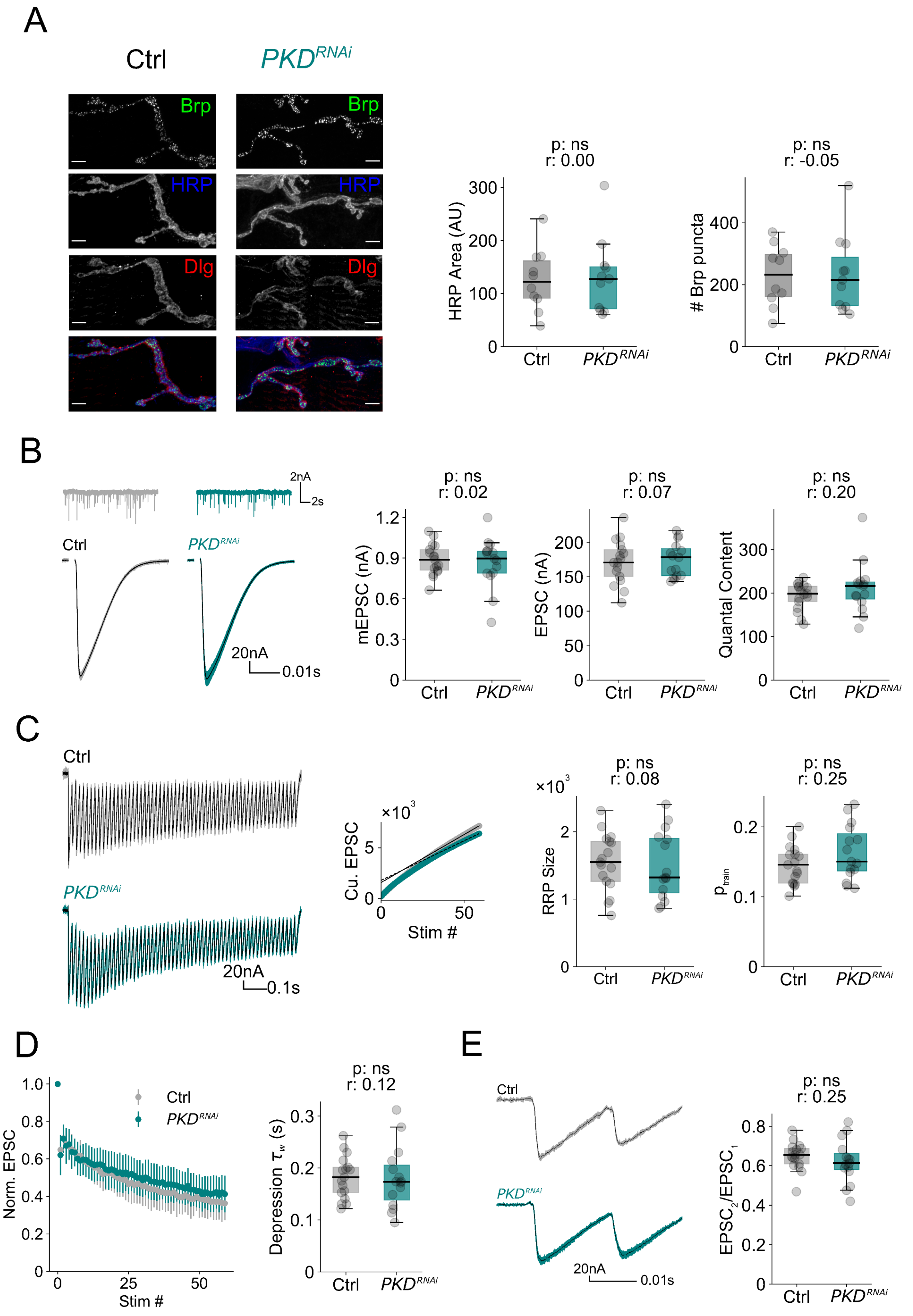

A
Ctrl
PKD RNAi
Brp
HRP
Dlg
p: ns
r: 0.00
HRP Area (AU)
p: ns
r: -0.05
### Brp puncta
B
2nA
2s
20nA
0.01s
mEPSC (nA)
p: ns
r: 0.02
EPSC (nA)
p: ns
r: 0.07
Quantal Content
p: ns
r: 0.20
C
0.1s
Cu. EPSC
Stim #
RRP Size
p: ns
r: 0.08
p train
p: ns
r: 0.25
D
Norm. EPSC
Depression τw (s)
p: ns
r: 0.12
E
EPSC2/EPSC1
p: ns
r: 0.25

**Fig. S6**. PKD knockdown does not affect NMJ morphology or baseline synaptic transmission. (**A**) Representative immunostainings for the active-zone marker Bruchpilot (Brp), neuronal membrane (HRP), and the postsynaptic reticulum (Dlg), along with quantification of synaptic (HRP) area and number of Brp puncta for NMJs without (Ctrl) or with ($PKD^{RNAi}$) PKD knockdown. Scale bar = 5 µm. n = 10 (Ctrl) vs. 11 ($PKD^{RNAi}$). (**B**) Representative traces (mEPSCs and EPSCs), mEPSP amplitudes, EPSC amplitudes and quantal content for NMJs without (Ctrl) or with ($PKD^{RNAi}$) PKD knockdown. (**C**) Representative EPSC trains (60 stimuli at 60 Hz frequency), linear fitting of last 10 cumulative EPSCs along with RRP size and release probability ($p_{train}$) estimated from cumulative EPSCs for NMJs without (Ctrl) or with ($PKD^{RNAi}$) PKD knockdown. (**D**) Normalized EPSCs, weighted depression time constant of normalized EPSC amplitudes ($\tau_w$), and (**E**) paired-pulse ratio ($EPSC_2/EPSC_1$) for NMJs without (Ctrl) or with ($PKD^{RNAi}$) PKD knockdown. n = 17 (Ctrl) vs. 15 ($PKD^{RNAi}$). $[Ca^{2+}]_e$ = 1.0 mM. Individual NMJs are shown as gray data points along with the box plot. p-value abbreviated as p and effect size abbreviated as r.
